# Lactate-responsive lysine lactylation in *Campylobacter* revealed by antibody-based detection of a conserved post-translational modification

**DOI:** 10.1101/2025.10.24.684354

**Authors:** Abdi Elmi, Brendan W Wren

## Abstract

Lysine lactylation (Kla) links lactate metabolism to protein regulation yet remains largely unexplored in bacteria. We used a Kla-specific rabbit polyclonal anti–ε-N-L-lactyl-lysine antibody to detect for this modification in *Campylobacter jejuni* (11168H, 81-176) and *Campylobacter coli* (M8). Immunoblotting of whole-cell lysates resolved discrete, strain-modulated bands under basal conditions and after physiologically relevant L-lactate pulses (4 or 16 mM). A conserved 42–45 kDa species was detected across strains, including in the absence of added lactate. Sample-load titrations revealed additional low-intensity bands, consistent with substoichiometric occupancy typical of a regulatory post-translational modification (PTM). Because *Campylobacter*, a member of the Epsilonproteobacteria, relies on amino-acid–driven, microaerophilic metabolism, we propose that endogenous and host-derived lactate generates lactyl donors (e.g., lactyl-CoA or lactoyl-glutathione) that enable Kla. We hypothesise that a CobB-like sirtuin mediates reversibility. To our knowledge, these data provide the first antibody-based evidence of Kla in *Campylobacter* and establish a framework for site-resolved mapping and functional tests that connect redox state and carbon flux to protein function, stress responses, and host interactions.

**Significance Statement:** Lactate, classically a metabolic end product, can modify lysine residues to regulate proteins, but this process is largely uncharted in bacteria. We examined *Campylobacter jejuni* and *Campylobacter coli* and detected lysine lactylation using a polyclonal antibody recognising **ε-**N-L-lactyl-lysine. Immunoblotting of whole-cell lysates revealed discrete, strain-modulated signals under basal conditions and after exposure to physiologically relevant L-lactate. A recurring 42–45 kDa band was present across strains, including in the absence of added lactate, and additional faint bands emerged with greater sample load, consistent with selective, low-occupancy modification. Because *Campylobacter* relies on amino-acid–driven metabolism under microaerophilic conditions, we suggest that endogenous and host-derived lactate generates activated lactyl donors (e.g., lactyl-CoA or lactoyl-glutathione) that support lysine lactylation, with a CobB-like sirtuin providing reversibility. These findings provide, to our knowledge, the first antibody-based evidence for lysine lactylation in *Campylobacter* and establish a basis for mapping sites and testing how metabolic state influences protein function, stress responses, and interactions with the hosts.

## Introduction

Lysine lactylation (Kla) has emerged as a mechanistic link between cellular metabolism and protein regulation, showing that lactate, a classical end product of glycolysis, can modify lysine residues and reshape enzyme activity and gene expression (Zong *et al*., 2025). In eukaryotes, including human macrophages, mouse embryonic stem cells and *Saccharomyces cerevisiae*, Kla couples intracellular lactate accumulation to transcriptional activation and chromatin remodelling via histone lysine modification (Zhang *et al*., 2019; Zhang *et al*., 2025). These observations recast lactate from a passive by-product to a dynamic signalling intermediate that links energetic state to proteome function (Wang *et al*., 2023; Zhao *et al*., 2024; Ren *et al*., 2025).

By contrast, investigation of similar systems in bacteria remains fragmentary. *Campylobacter jejuni* and *Campylobacter coli* are major causes of acute bacterial gastroenteritis and occupy a distinctive metabolic niche (Elmi *et al*., 2020; Omole *et al*., 2024). Unlike enteric Gammaproteobacteria such as *Escherichia coli* and *Salmonella enterica*, *Campylobacter* lacks key glycolytic enzymes, including glucokinase (*glk*) and phosphofructokinase (*pfkA*), and therefore does not use a canonical Embden–Meyerhof–Parnas pathway (Parkhill *et al*., 2000). Excluding a minority of strains harbouring an L-fucose utilisation locus (*fuc*, *cj0480c–cj0490*) that permits limited fucose metabolism, *C. jejuni* relies on a condensed, amino-acid–driven network in which serine, aspartate and glutamate catabolism converge on pyruvate (Stahl *et al*., 2012). Under microaerophilic conditions, pyruvate fuels the tricarboxylic acid (TCA) cycle to sustain oxidative respiration and biosynthesis, an evolutionarily streamlined, energy-efficient programme that contrasts with the fermentative strategies of classical enteric pathogens (Stahl *et al*., 2011; Hofreuter, 2014; Burnham & Hendrixson, 2018).

In oxygen-limited, host-like atmospheres (5% O₂, 10% CO₂, 85% N₂), *C. jejuni* may generate small amounts of L-lactate from pyruvate. Because it lacks a canonical cytoplasmic NADH-dependent LDH and relies on quinone-linked periplasmic LDHs that oxidise lactate to pyruvate, any reverse flux is likely minor, context-dependent and mechanistically unresolved *in vivo*. In primary hosts (e.g., the chicken caecum) and incidental human intestinal niches, lactate can accumulate to millimolar levels (approximately 4–30 mM), positioning it as both a carbon source and a potential signalling molecule. Under such conditions, formation of activated lactyl donors, putatively lactyl-CoA and/or lactoyl-glutathione could drive ε-lysine lactylation on selected proteins via enzymatic or non-enzymatic routes, altering side-chain charge/polarity and thereby modulating conformation, stability and interaction networks.

Within this biochemical and ecological context, *Campylobacter* provides a tractable model for testing how redox-active metabolites feed into post-translational regulation. Its reliance on amino-acid catabolism, together with exposure to host-derived lactate in primary avian and incidental mammalian hosts, supplies both the metabolic precursors and environmental cues necessary for Kla, whether enzymatic or non-enzymatic. A mechanism in which pyruvate reduction and bidirectional respiration with host lactate tunes proteome composition would explain how organisms lacking classical glucose-based glycolysis dynamically adjust energy flux, stress responses and virulence traits.

Here, we investigate the hypothesis that Kla occurs in *Campylobacter* as a redox-sensitive modification arising from amino-acid catabolism and exogenous lactate exposure. Using the well-characterised wild-type strains *C. jejuni* 11168H and 81-176 and a wild-type *C. coli* isolate, we pulsed cultures with physiologically relevant concentrations of sodium L-lactate and probed lysates by immunoblot with a validated polyclonal anti–ε-N-L-lactyl-lysine antibody. Distinct immunoreactive bands were detected under both basal and lactate-enriched conditions, demonstrating that *Campylobacter* proteins undergo Kla in response to metabolic state. Because Kla relies on donors such as lactyl-CoA or lactoyl-glutathione, these data indicate that *Campylobacter* couples host- and environment-derived lactate to proteomic regulation. While comprehensive site mapping and structural analyses remain to be undertaken, our antibody-based detection provides, to our knowledge, the first evidence of Kla in *Campylobacter*, establishing a foundation for investigating metabolic–regulatory coupling in microaerophilic pathogens and suggesting that, at the host–pathogen interface, lactate acts not only as a carbon source but also as a signalling molecule that modulates *Campylobacter* redox physiology, adaptive behaviour, and virulence potential.

## Methods

### Bacterial strains and growth conditions

*Campylobacter jejuni* strains 11168H and 81-176, and a newly isolated wild-type *Campylobacter coli* strain M8, were maintained on Columbia blood agar (CBA; Oxoid, CM0331) at 37 °C under microaerophilic conditions (85% N₂, 10% CO₂, 5% O₂). Single colonies from first-passage plates were used to inoculate 10–15 mL Brucella broth (BB; Oxoid, CM0169) and were incubated for 12 h at 37 °C in the same atmosphere with gentle shaking (75 rpm). Cultures were diluted into pre-equilibrated BB to an initial OD600 ≈ 0.10 and grown in a Don Whitley microaerophilic workstation to early exponential phase (∼2–3 h) or mid-exponential phase (∼4–5 h), OD600 ranges (e.g., 0.2–0.3 / 0.4–0.6) as indicated.

### Lactate pulsing and control treatments

At the indicated growth phase, cultures were pulsed with freshly prepared sodium L-lactate (Sigma-Aldrich, L7022) to final concentrations of 4 mM or 16 mM. Control cultures received an equal volume of sterile BB or were left untreated. Following addition, cells were incubated for 60–90 min at 37 °C under microaerophilic conditions.

### Cell harvesting, lysis, and protein quantification

Cells were collected by centrifugation (4,000 × g, 25 min, 4 °C), washed once with ice-cold PBS, and resuspended in ice-cold RIPA buffer (50 mM Tris-HCl, pH 7.5; 150 mM NaCl; 1% NP-40; 0.5% sodium deoxycholate; 0.1% SDS; 1 mM EDTA) supplemented with protease inhibitors (Roche, 11836153001). Lysates were clarified (14,000 × g, 15–20 min, 4 °C). Total protein was measured with the Qubit™ Broad Range Protein Assay (Thermo Fisher, Q33211). Samples were normalised to 0.05–1.5 µg µL⁻¹, mixed with 4× Laemmli buffer containing 100 mM DTT, and heated at 65 °C for 5 min prior to SDS–PAGE.

### SDS–PAGE and Western blot detection of lysine lactylation

Equal amounts of total protein (from samples normalised to 0.05–1.5 µg µL⁻¹) were resolved by SDS–PAGE (Invitrogen, UK) and dry-transferred using an iBlot 3 system (Invitrogen, IB33001). Membranes were blocked for 1 h in 5% (w/v) BSA in PBST (NaCl, 8 g; KCl, 0.2 g; Na₂HPO₄, 1.44 g; KH₂PO₄, 0.24 g; 0.1% Tween-20; pH 7.6 per litre) and incubated overnight at 4 °C with rabbit polyclonal anti–ε-N-L-lactyl-lysine antibody (1:1,000; Fisher, 17219403) in 5% (w/v) BSA in PBST. After five PBST washes, membranes were incubated for 1 h at room temperature with IRDye 800CW–conjugated anti-rabbit IgG (1:5,000; LI-COR, 926-32211), protected from light. Near-infrared fluorescence was acquired on an Odyssey M imaging system (LI-COR) under non-saturating conditions. Loading was verified by total-protein fluorescence of the same membrane prior to antibody probing.

### Image quantification and analysis

Fluorescent immunoblots were analysed in LI-COR Image Studio Lite v5.2. Bands were quantified within the linear range after local background subtraction. Integrated intensities were normalised to total-protein fluorescence from the same membrane to correct for lane-to-lane variation. Data were exported as relative fluorescence units (RFU) for comparison. Figures were prepared in FIJI (ImageJ v1.54) without additional contrast adjustment.

### Statistical Analysis

Statistical analyses were performed in GraphPad Prism (v10.6.1; GraphPad Software).

## Results

### Lysine lactylation is selective, sub-stoichiometric, and linked to metabolic flux

We profiled Kla in *C. jejuni* 11168H, *C. jejuni* 81-176, and a wild-type *C. coli* isolate (strain M8) under microaerophilic conditions (Fig. 1A). Cultures were harvested at early to mid-log (OD600 ≈ 0.3) shortly after pulsed with 4 mM or 16 mM sodium L-lactate. These concentrations approximate exposure in avian and mammalian gut niches. Growth and total protein were quantified by OD600 and Qubit, and samples were normalised across conditions (Fig. 1B–C).

**Figure 1.**
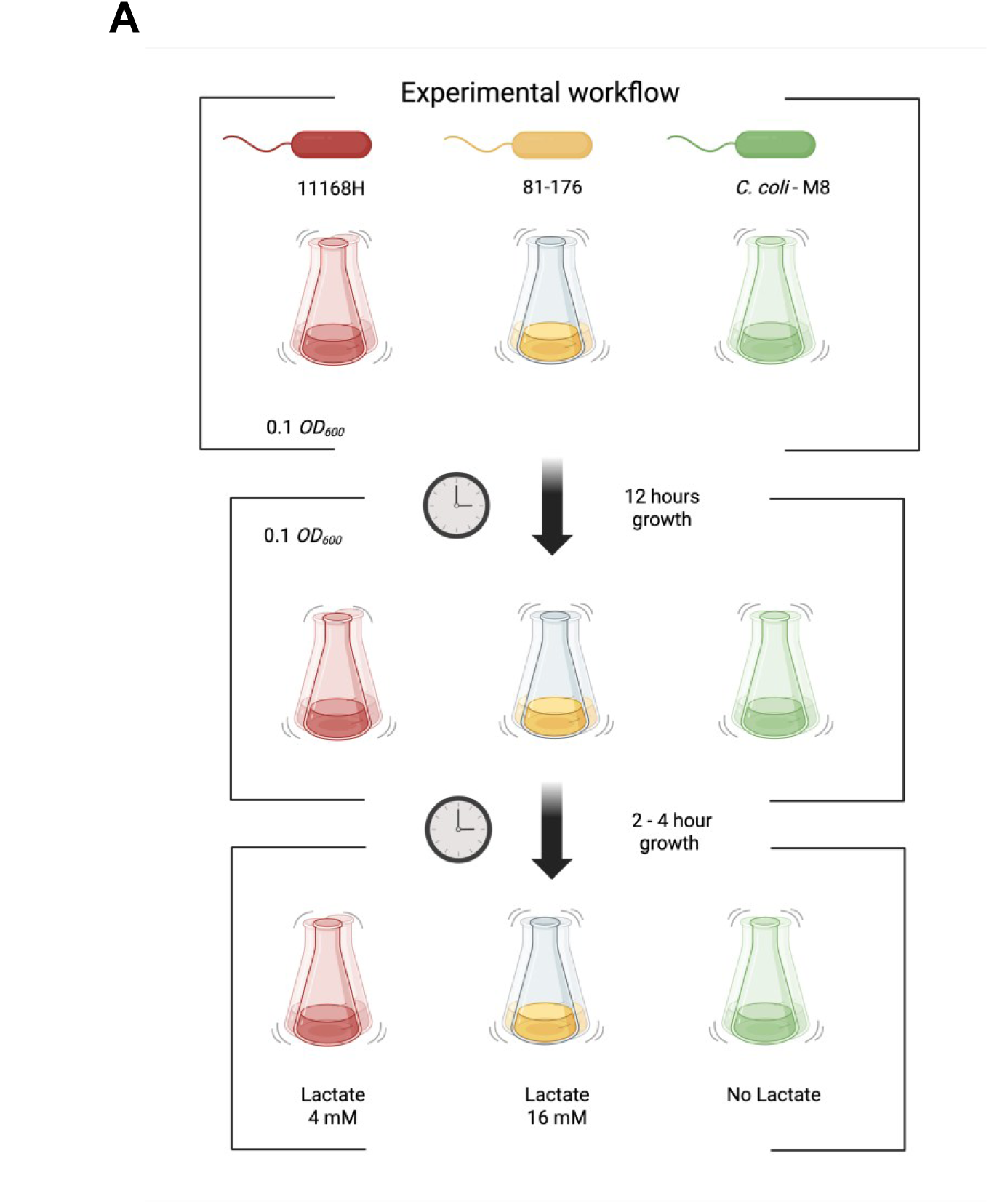
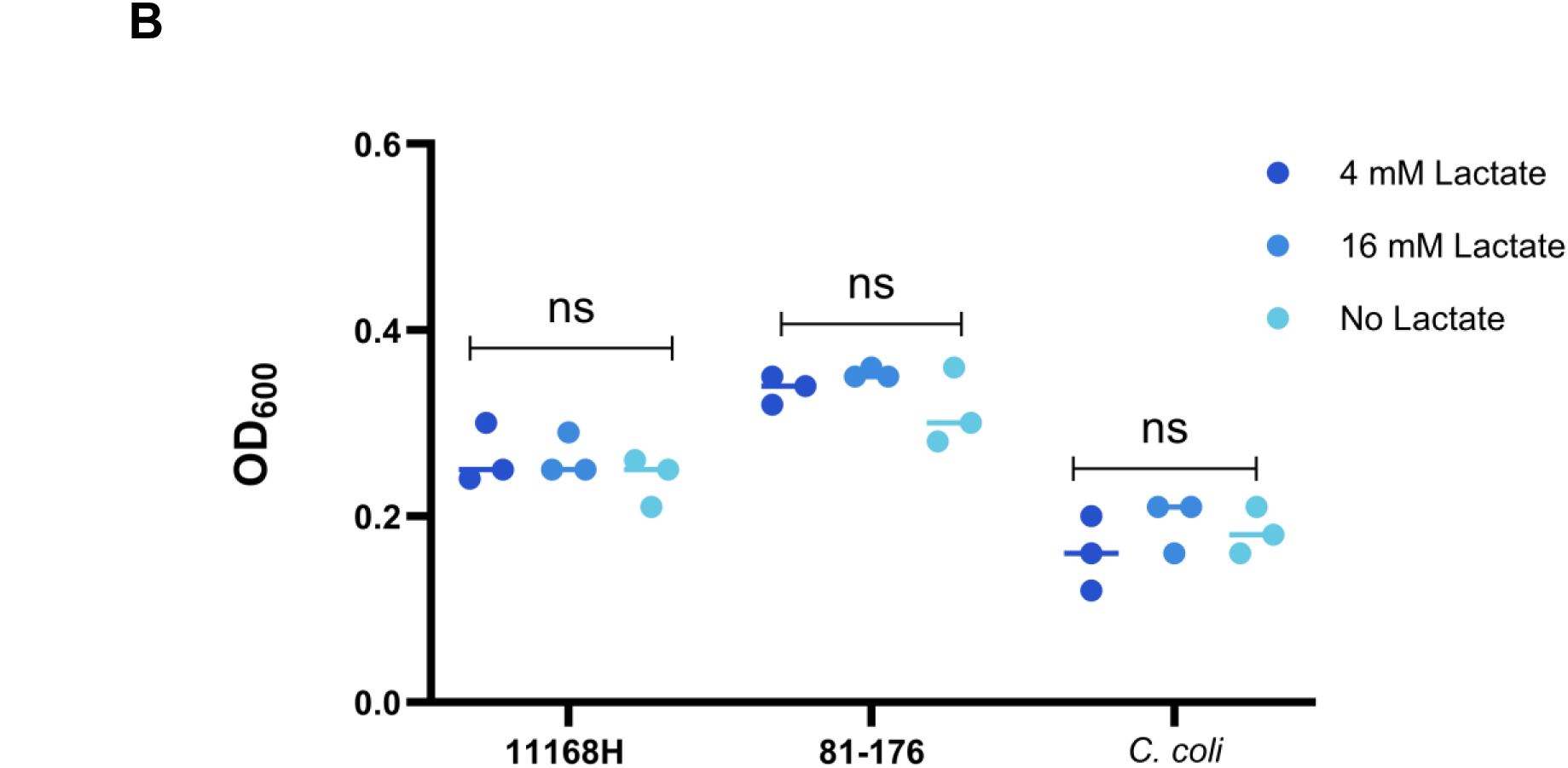
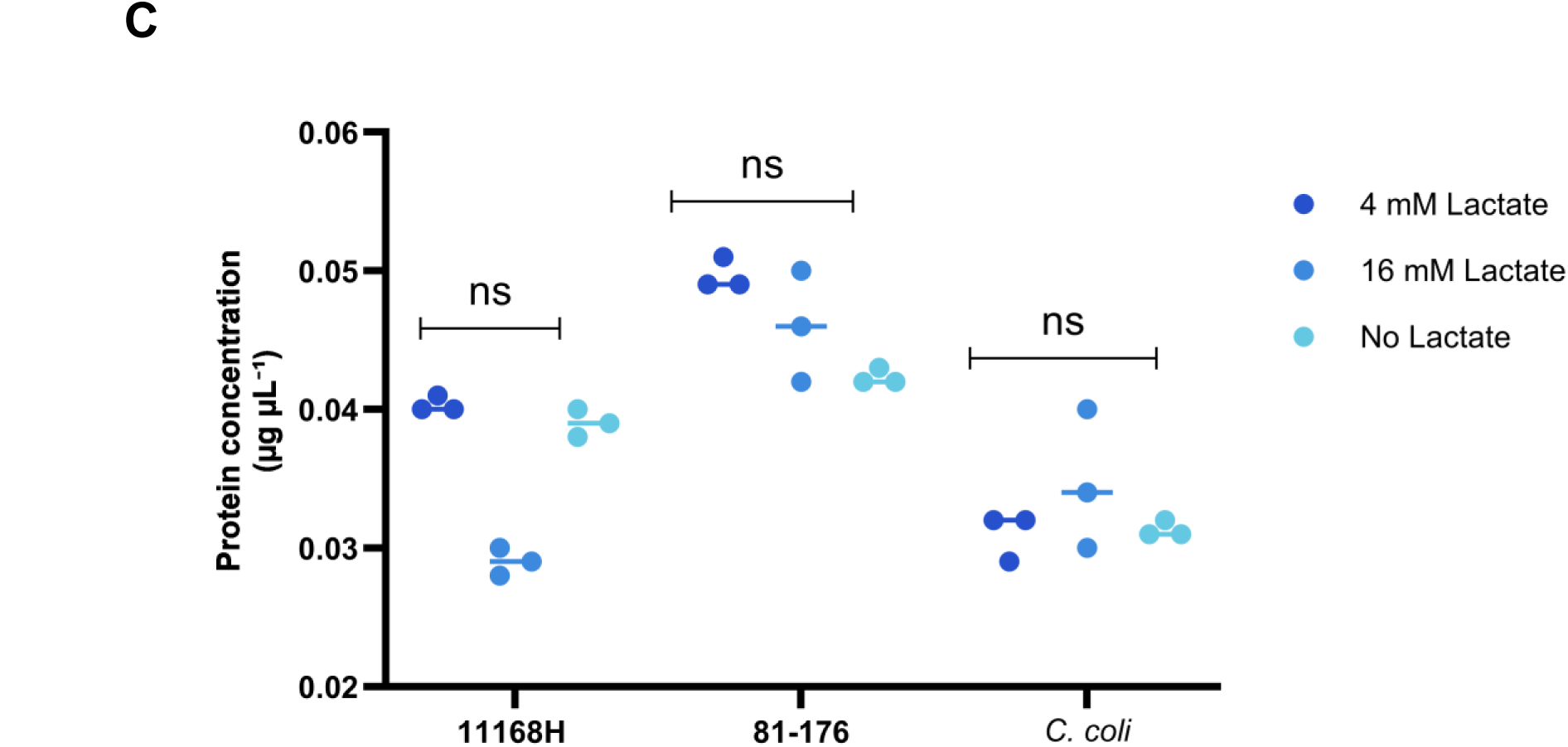
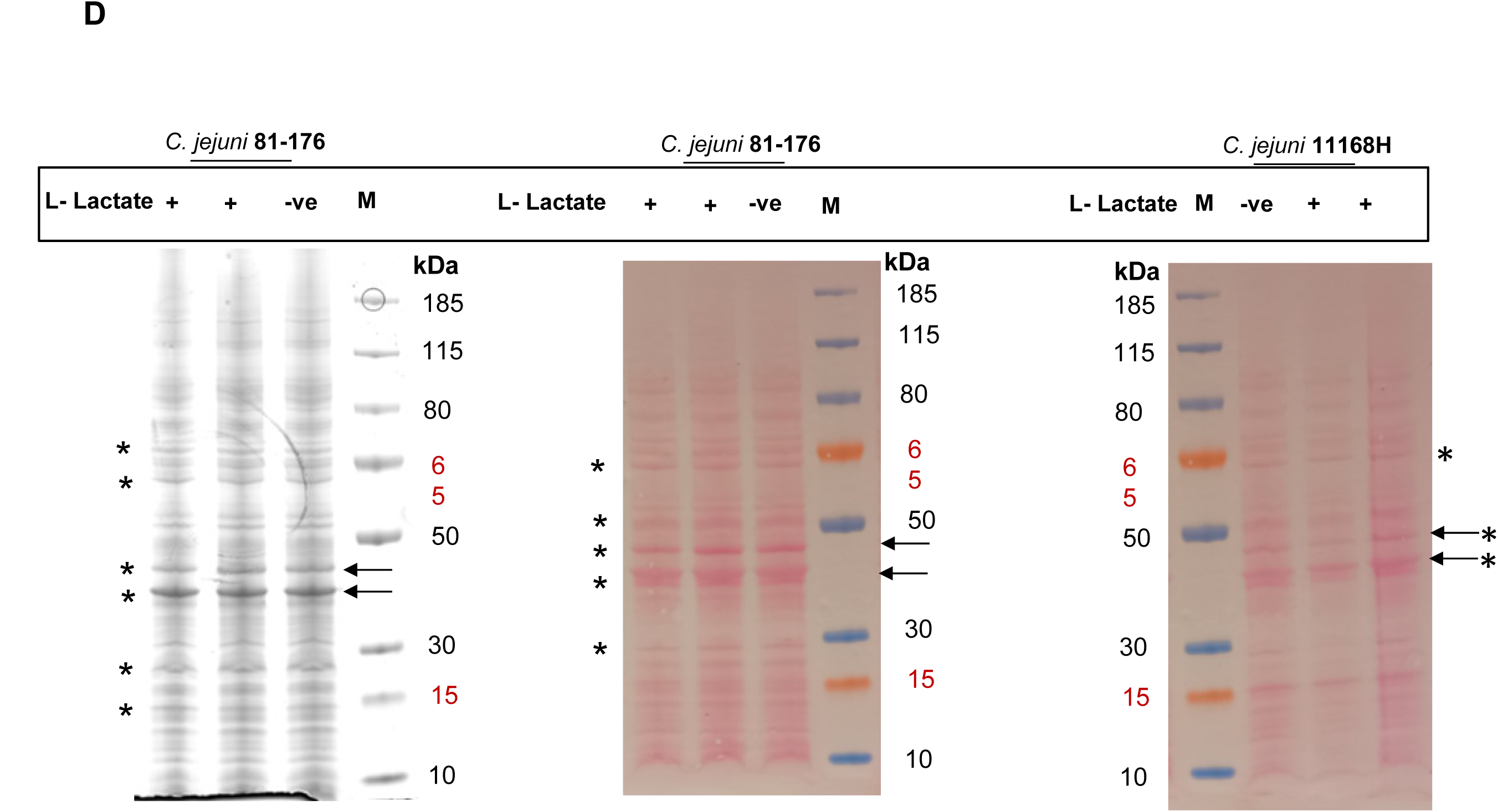
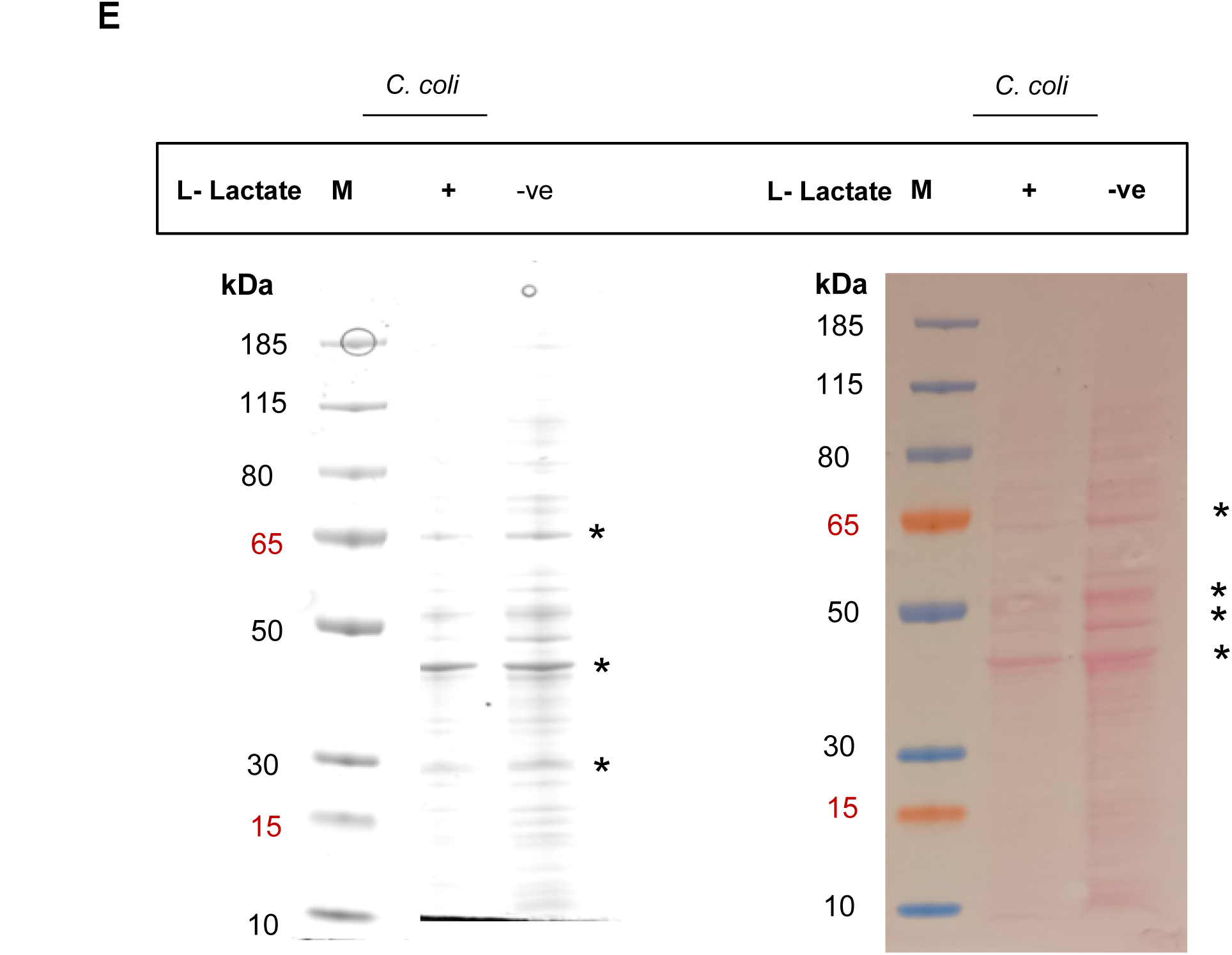
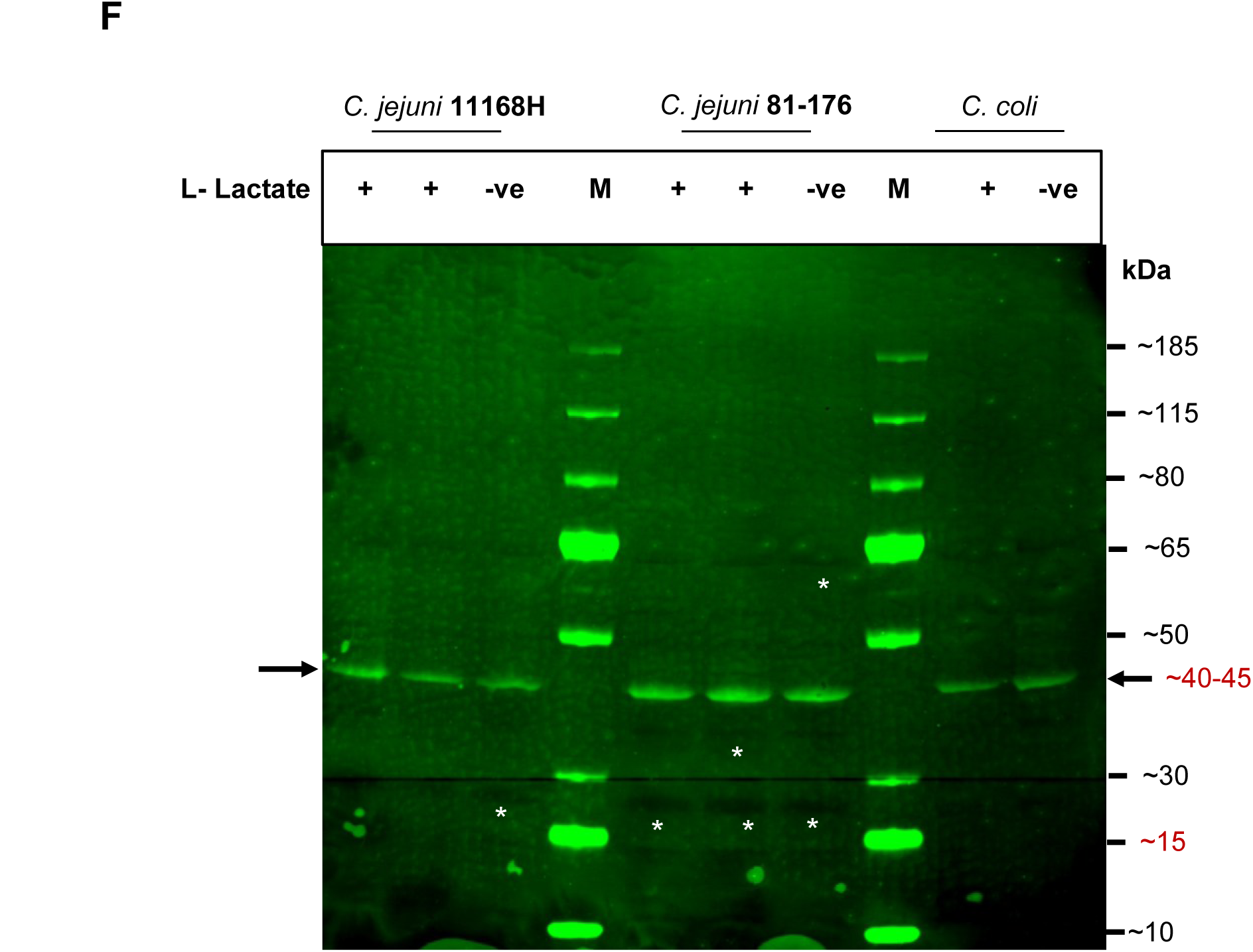
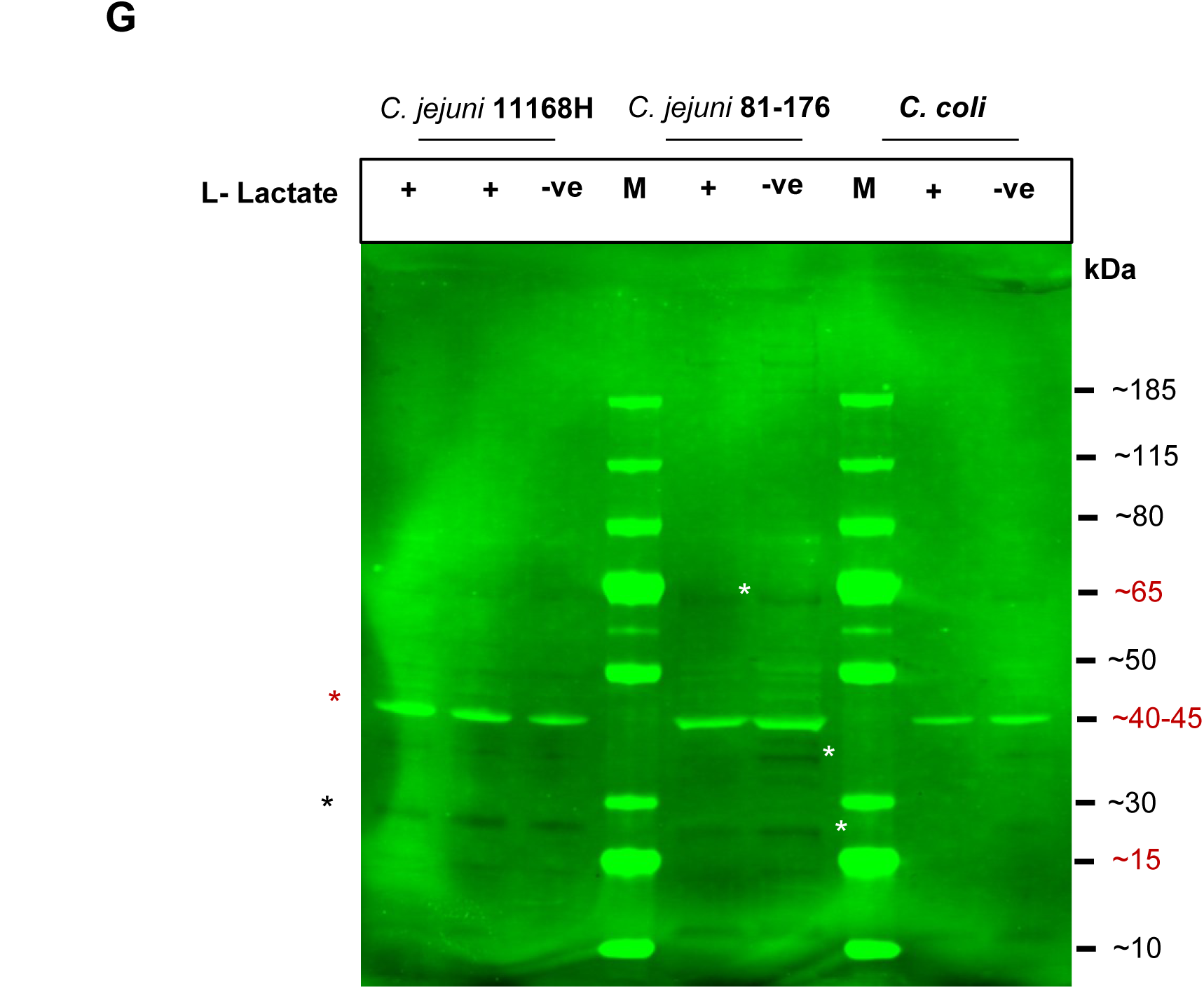
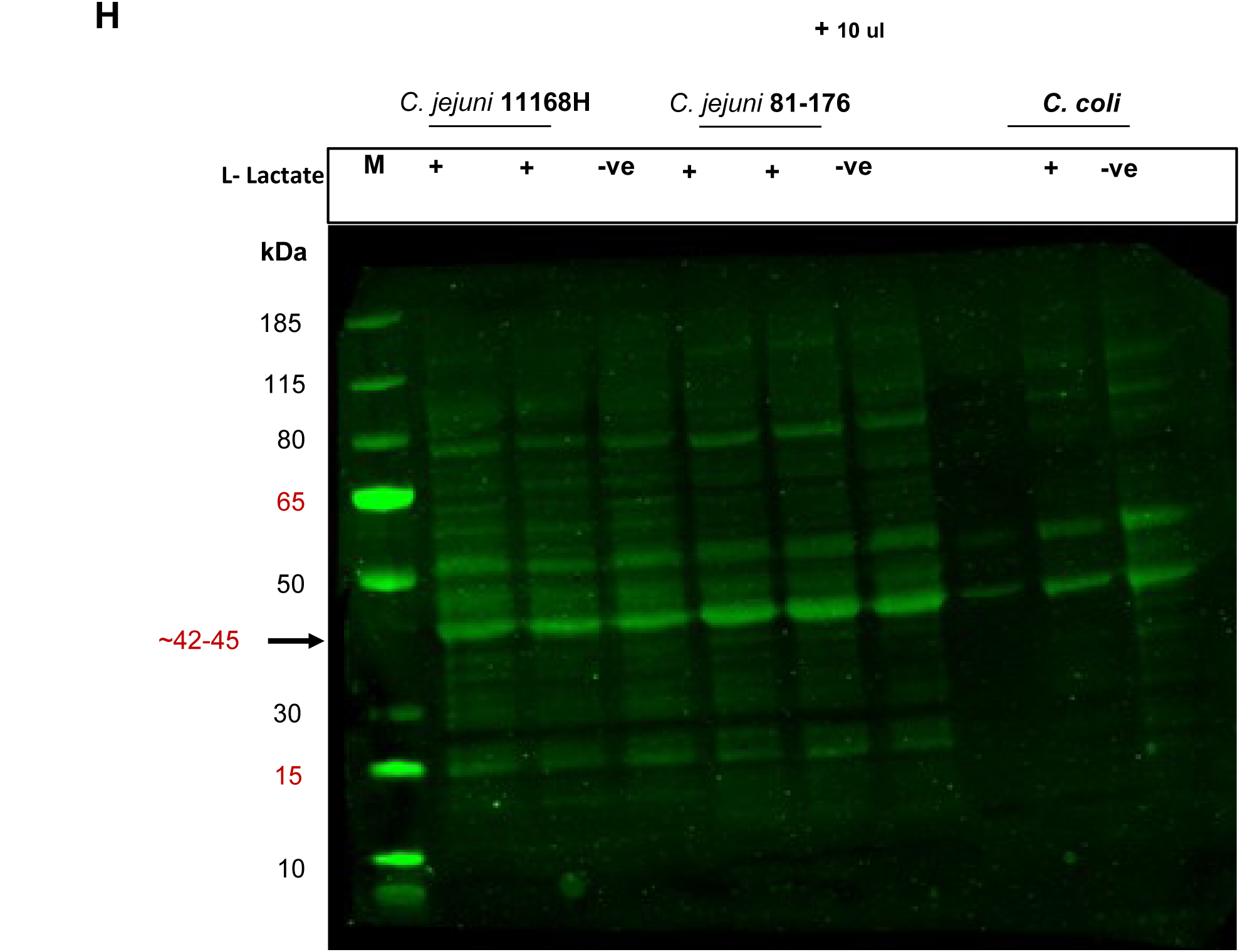
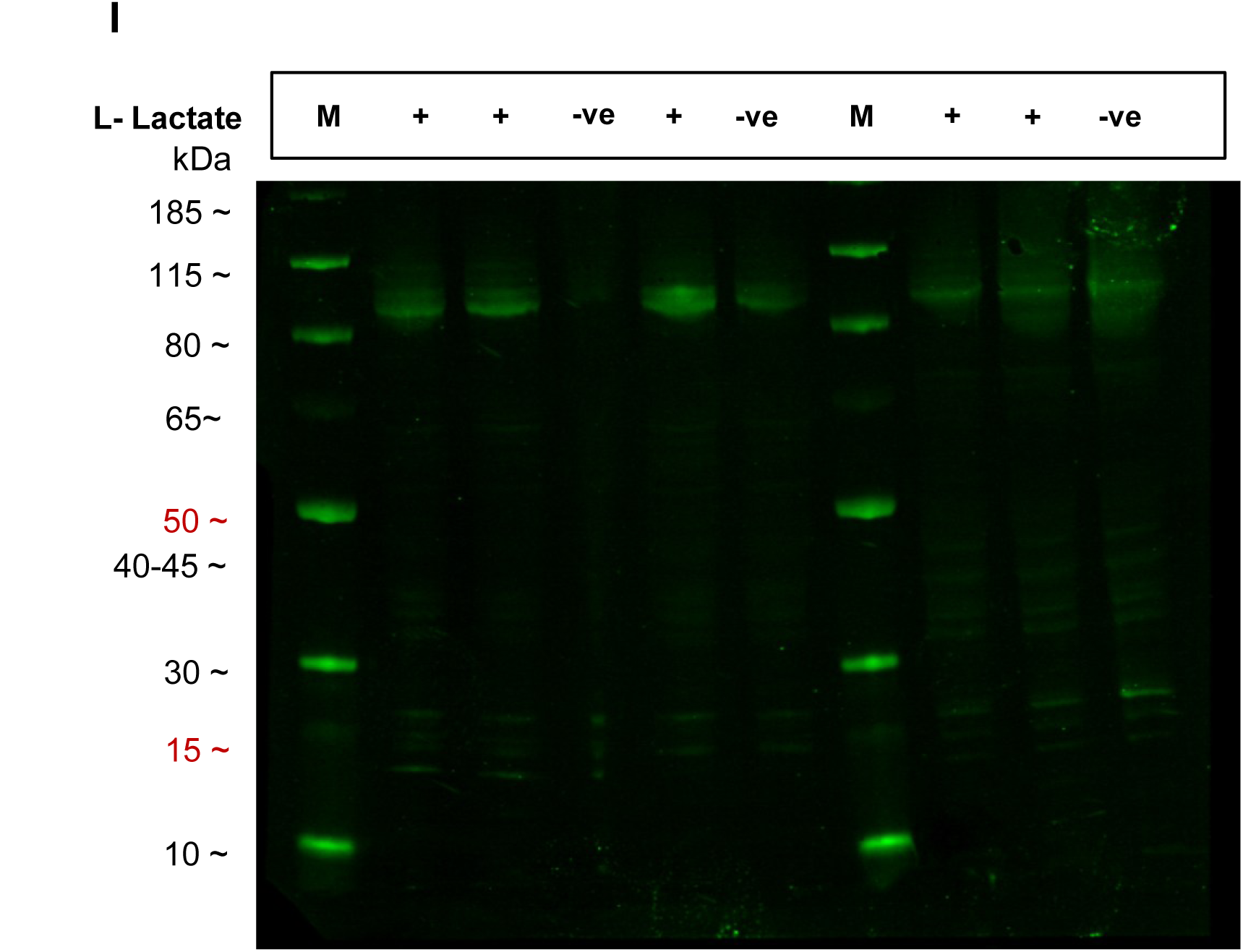
Selective Kla during early–mid-log (OD600 ≈ 0.3) reveals specificity beyond protein abundance. **(A)** Experimental workflow. Cultures were pulsed with 4 or 16 mM L-lactate, rapidly chilled to 4 °C, and lysed in RIPA buffer. **(B)** Total protein. Qubit measurements for *C. jejuni* 11168H, 81-176, and *C. coli* strains (± lactate biological duplicates), used for loading normalisation. **(C)** Bacterial growth at sampling. Within-strain OD600 at early–mid log (≈0.3) was indistinguishable across conditions (no significant differences), confirming equivalent cellular biomass per lane. **(D–E)** Loading/transfer verification. Imperial Protein Stain and Ponceau S confirmed loading/transfer; some abundant bands (asterisks) lack Kla signal. **(F–G)** Kla immunoblot. Discrete bands detected under basal and lactate-stimulated conditions; a constitutive ∼42–45 kDa species is present without exogenous lactate. **(H)** Input titration (10→30 µL) reveals additional faint Kla bands, indicating sub-stoichiometric occupancy; strain-specific differences suggest possible regulation by redox flux and/or delactylases. **(I)** Secondary-only control for antibody specificity. Membrane incubated with IRDye 800CW goat anti-rabbit IgG alone produced only two faint, high-molecular-weight bands across all samples.

Total-protein staining of the same membrane confirmed matched loading; minor, treatment-dependent intensity differences were within the validated linear range (Fig. 1D–E), consistent with strain-specific physiology rather than technical variance. Immunoblotting with a validated anti–ε-N-L-lactyl-lysine antibody revealed distinct, strain-specific Kla profiles that did not correlate with total protein abundance (Fig. 1F–G). A reproducible band at ∼42–45 kDa was detected across all strains and conditions, including in the absence of added lactate, indicating constitutive lactylation of at least one protein. Increasing lysate load from 10 to 30 µL revealed additional low-intensity bands (Fig. 1H), consistent with sub-stoichiometric occupancy typical of regulatory PTMs. Control blots for antibody specificity, incubated with secondary antibody only, (IRDye 800CW goat anti-rabbit IgG) showed just two faint, high–molecular-weight bands across all samples (Fig. 1I). At both early mid-log and mid-log, total protein remained within a narrow range, indicating that variation in Kla intensity reflects intrinsic regulation rather than biomass or extraction efficiency. Together, these data establish Kla in *Campylobacter* as a selective and regulated modification, sub-stoichiometric, and responsive to metabolic flux.

### Lactate-independent Kla persists at mid-log and is strain-modulated

At mid-log (OD600 ≈ 0.4–0.55), *C. jejuni* 11168H, *C. jejuni* 81-176, and *C. coli* M8 were cultured with or without 4 or 16 mM L-lactate. Growth and Qubit measurements were consistent across conditions, supporting normalisation across conditions (Fig. 2A–B). SDS– PAGE gels stained with Imperial Protein Stains showed modest, reproducible differences among strains; notably, 11168H without lactate exhibited slightly reduced staining relative to its Qubit-determined load, mirrored by a fainter Kla signal on the corresponding immunoblot (Fig. 2C–D). Immunoblotting detected a robust Kla band at ∼42–45 kDa. across all strains, independent of exogenous lactate, indicating constitutive lactylation. This plausibly supported by endogenous lactate arising from peptide catabolism and serine- or pyruvate-derived flux (cf. Fig. 3A–D). The persistence of this species from early mid-log to mid-log supports a conserved, metabolically coupled lactylation process intrinsic to *Campylobacter’s* central metabolism.

**Figure 2.**
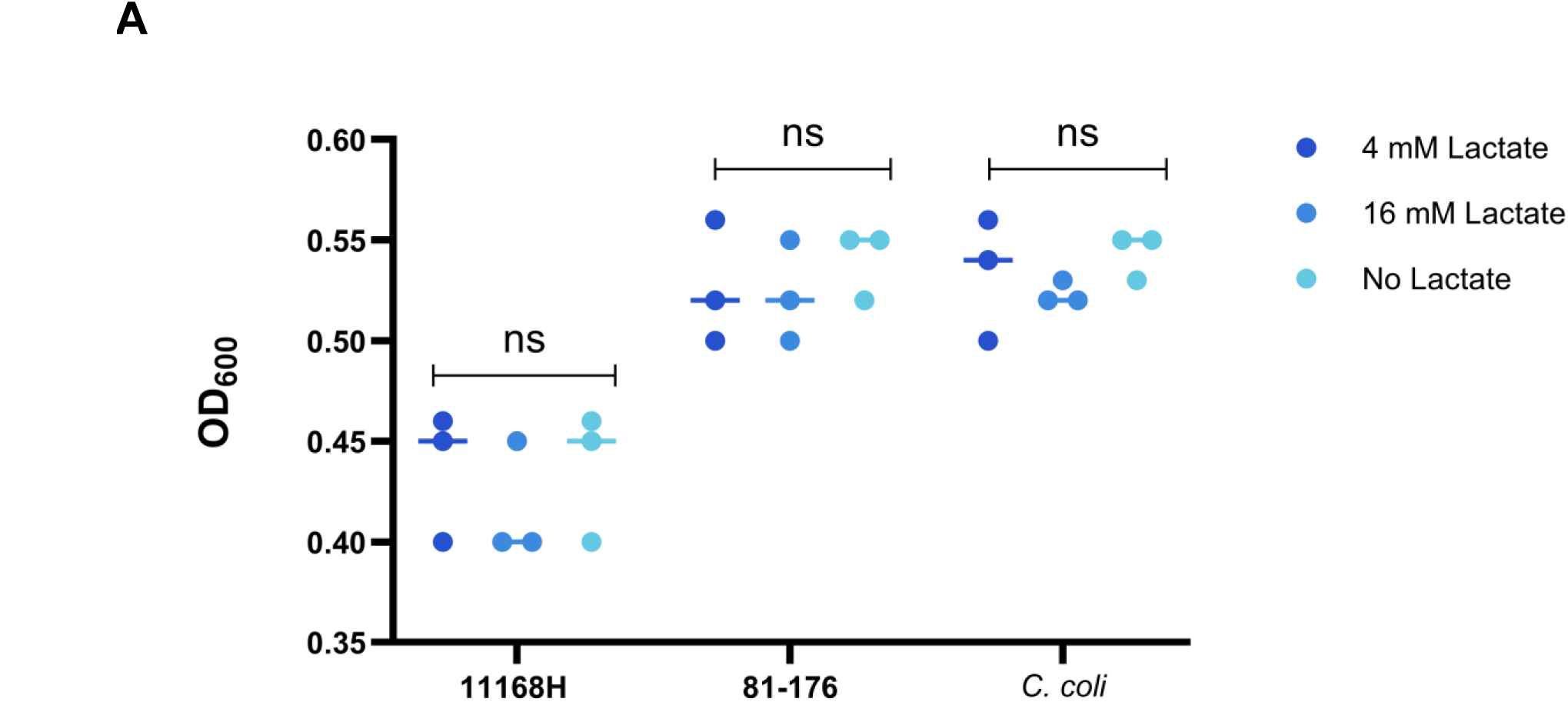
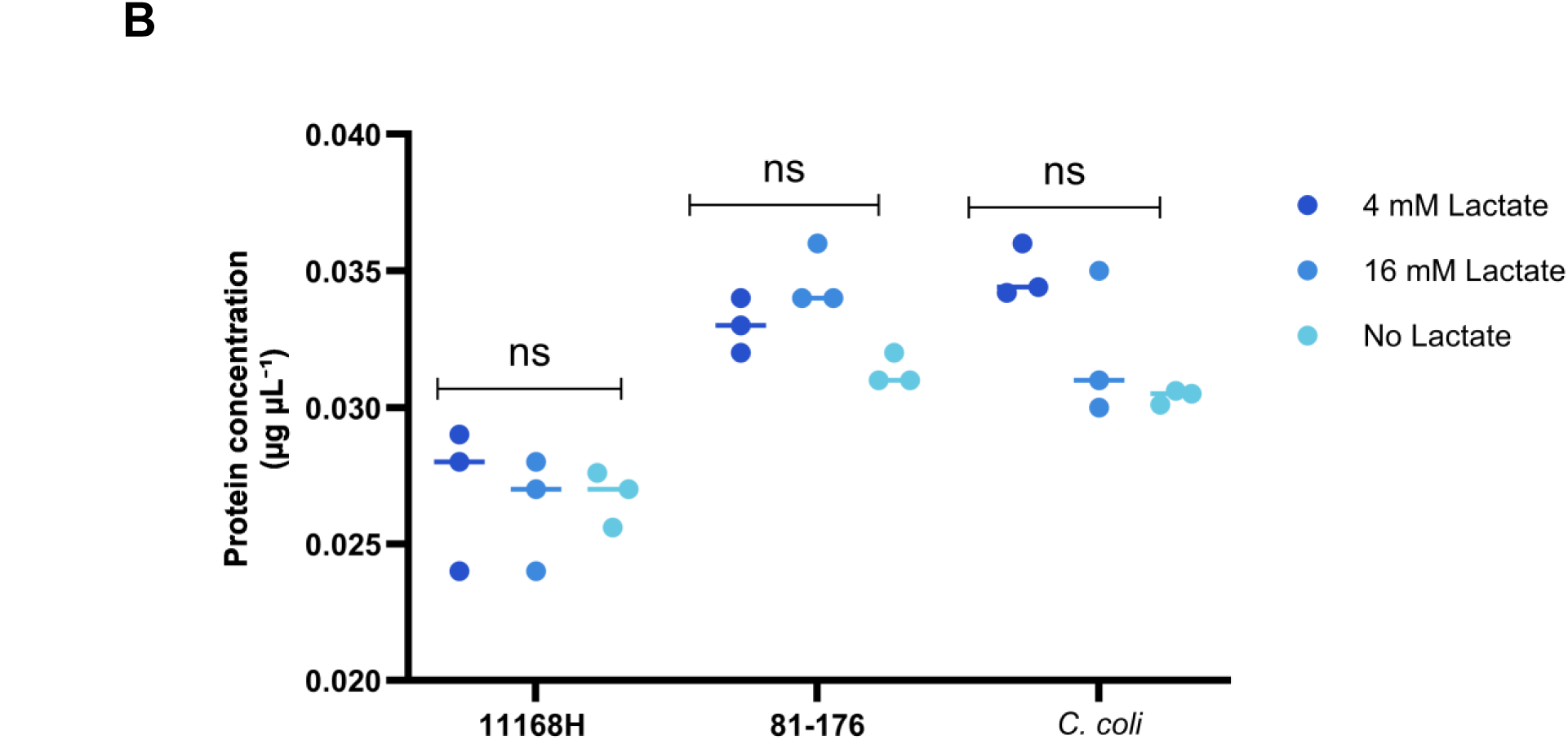
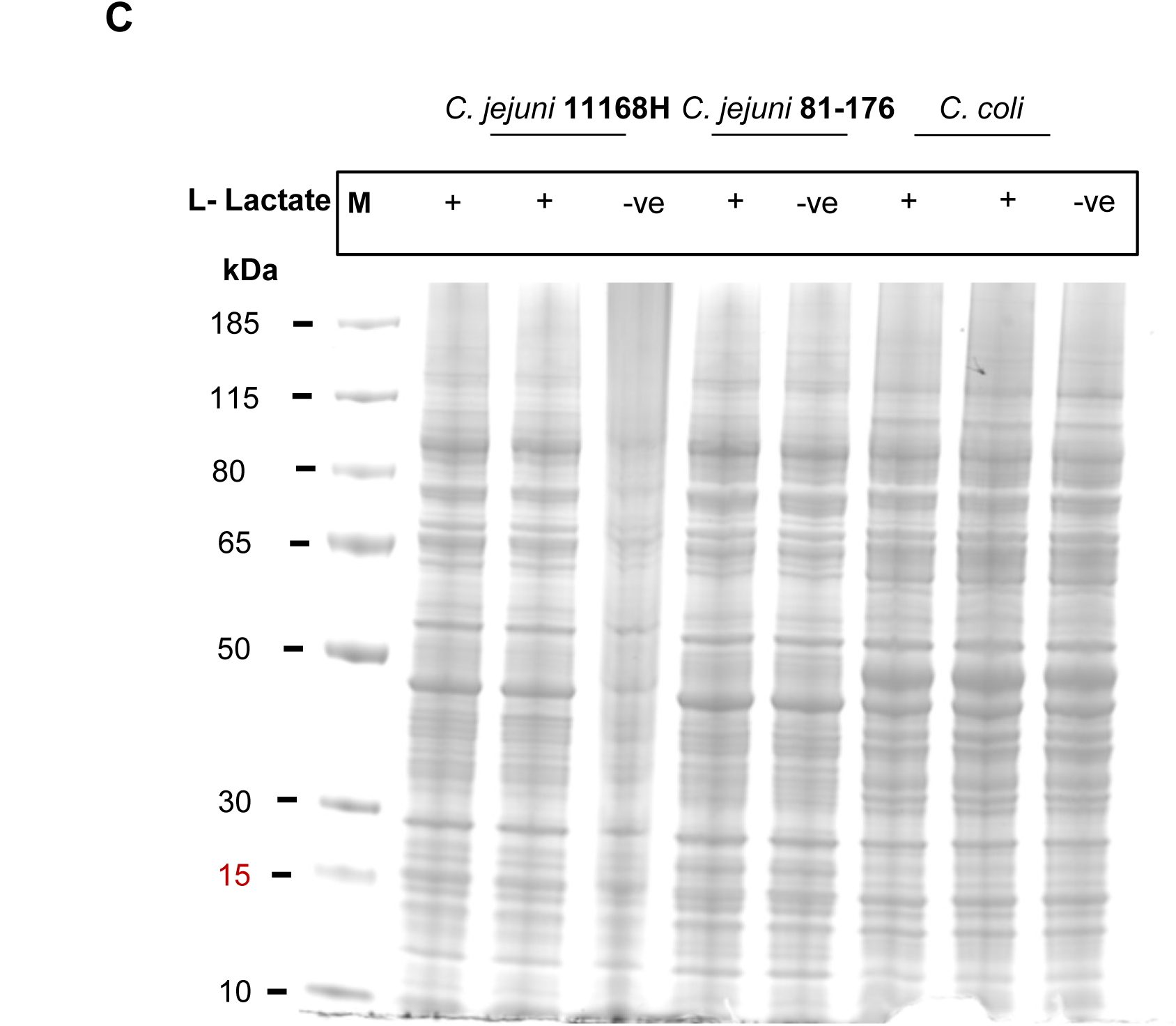
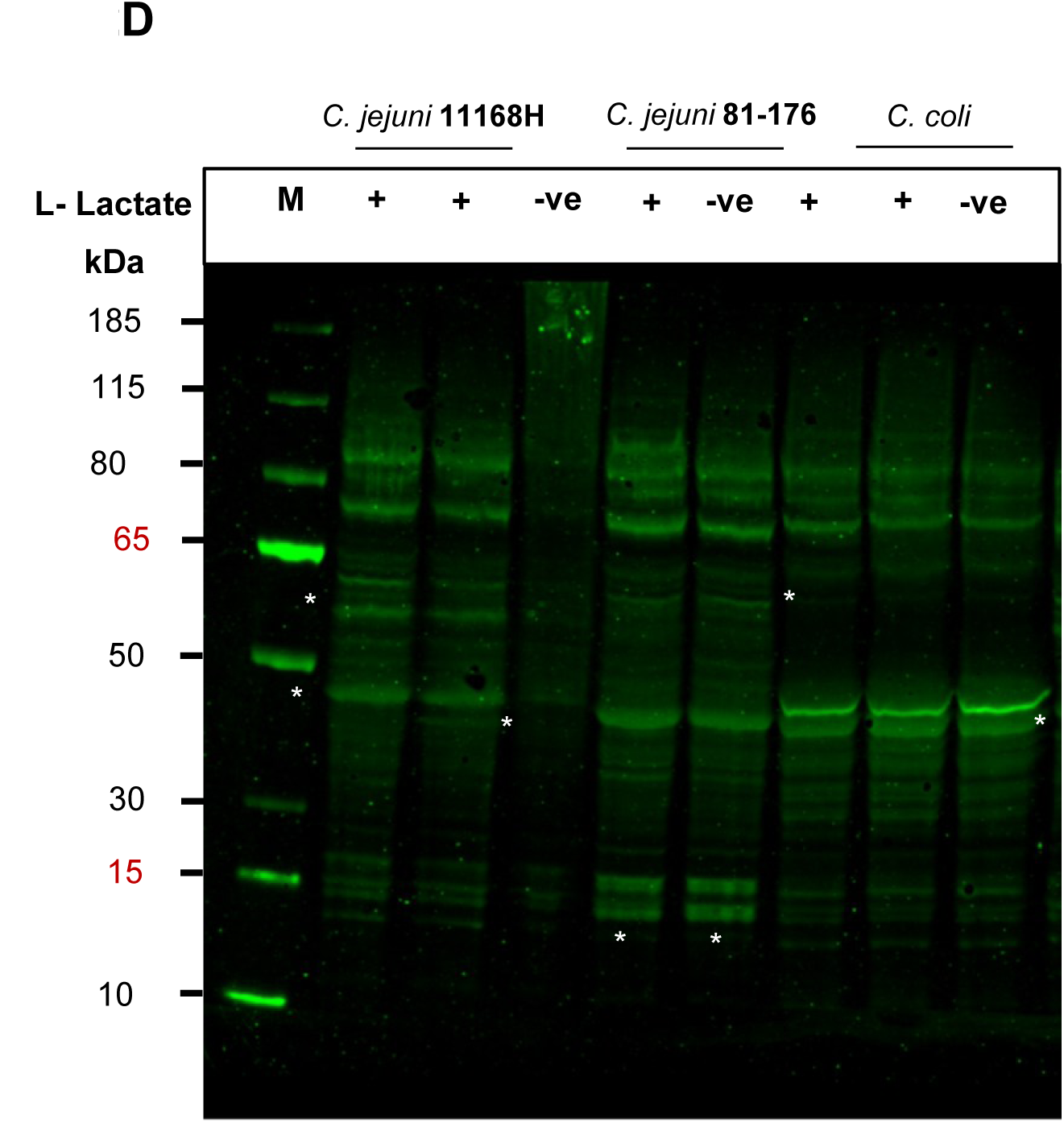
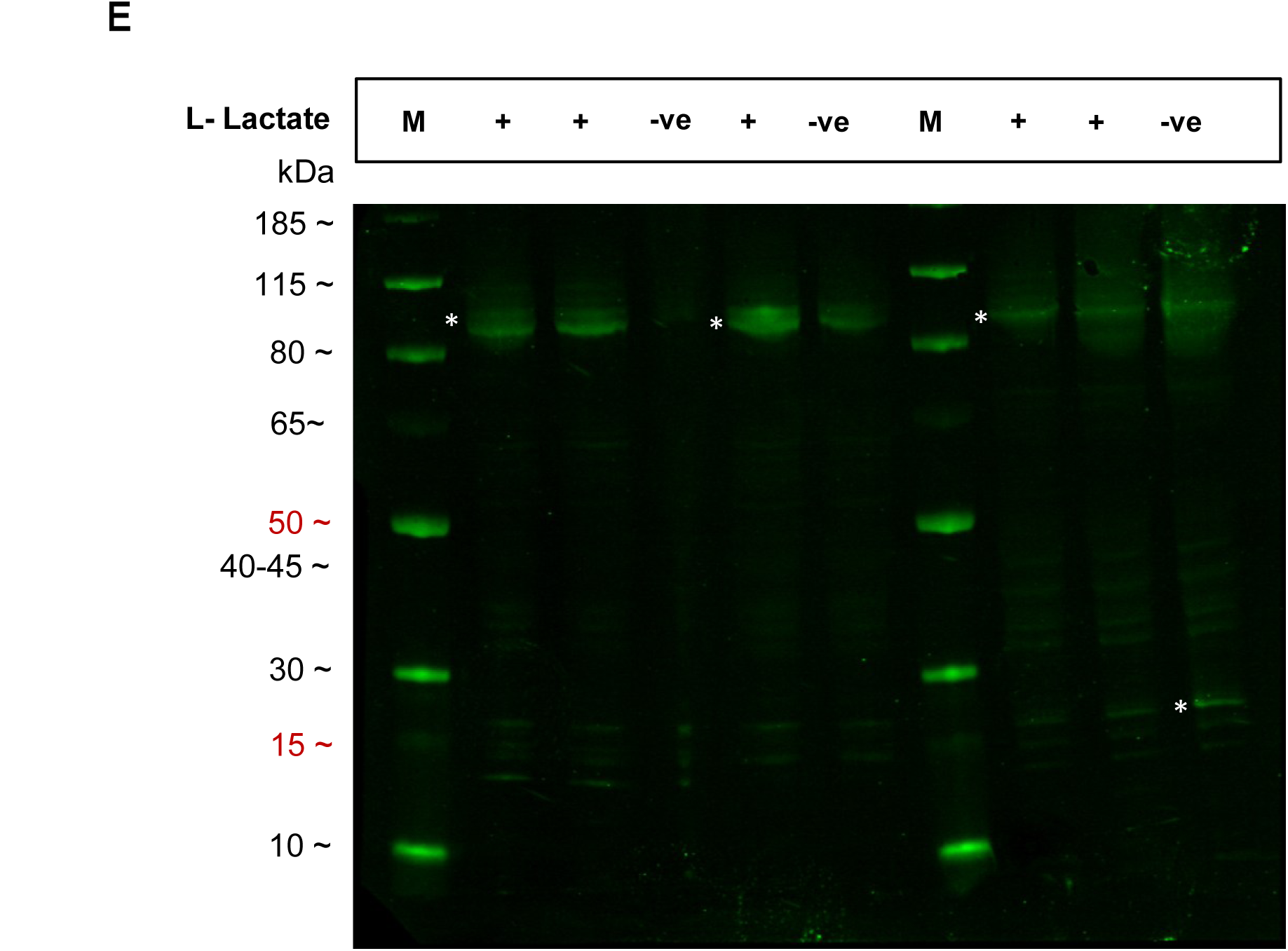
Kla expression during mid-log growth in *Campylobacter*. **(A)** Bacterial growth at sampling. Within-strain OD600 at early–mid log (≈0.5) was indistinguishable across conditions (no significant differences), confirming equivalent cellular biomass per lane. **(B)** Total protein. Qubit measurements for *C. jejuni* 11168H, 81-176, and *C. coli* strains ± lactate (duplicates) show comparable yields. **(C)** Loading verification. Imperial staining confirms loading; abundant proteins (Imperial) need not be Kla-positive. **(D)** Kla immunoblot. A prominent ∼42–45 kDa species is detected across strains and conditions; additional bands vary by strain, consistent with a regulated, redox-linked modification. **(E)** Control validation of anti-Kla specificity. Membranes containing increasing titrations of whole-cell incubated only with secondary IRDye 800CW goat anti-rabbit IgG. The absence of distinct signals beyond two faint, high–molecular-weight bands confirms that all Kla bands detected in experimental blots are primary–antibody dependent.

**Figure 3.**
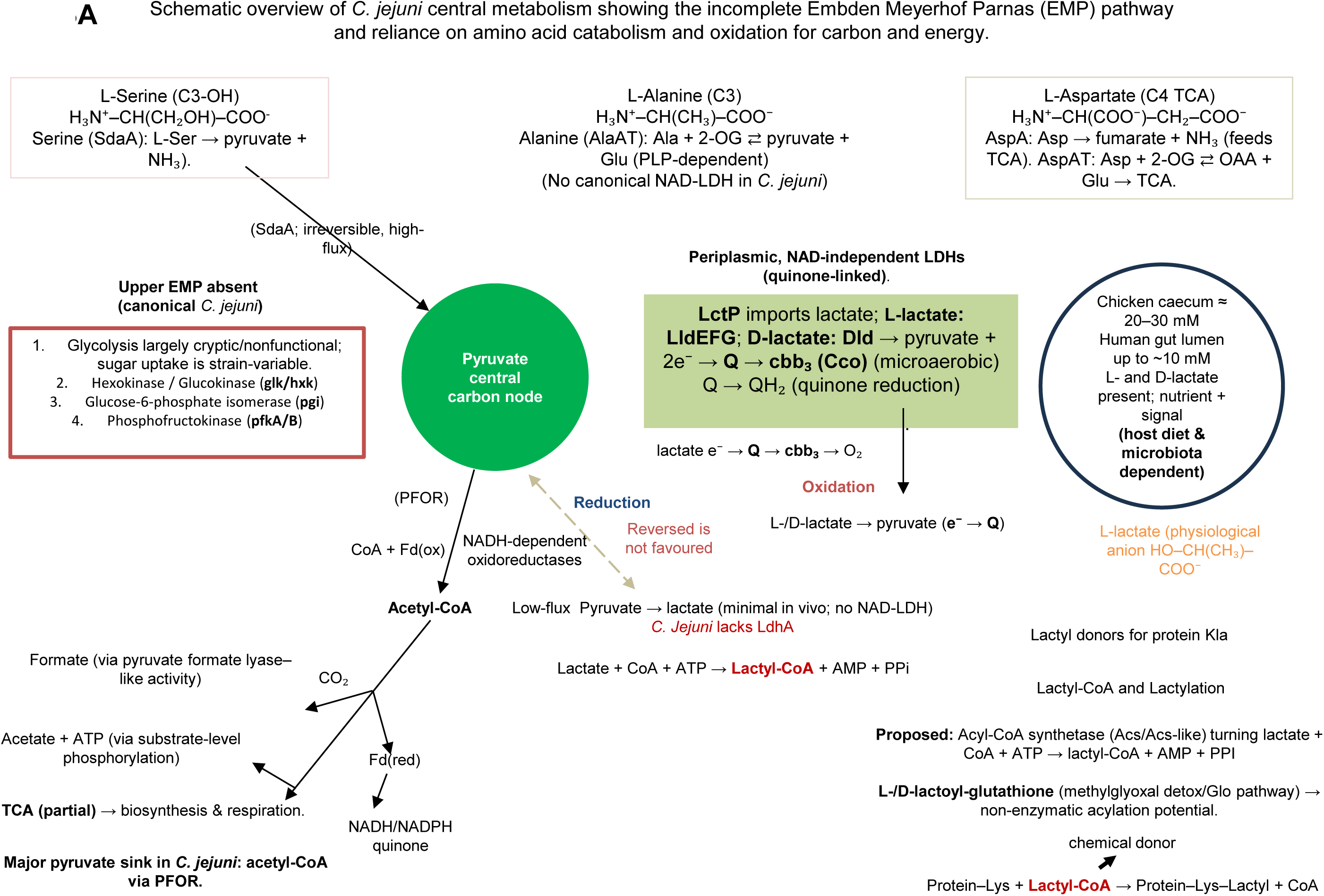
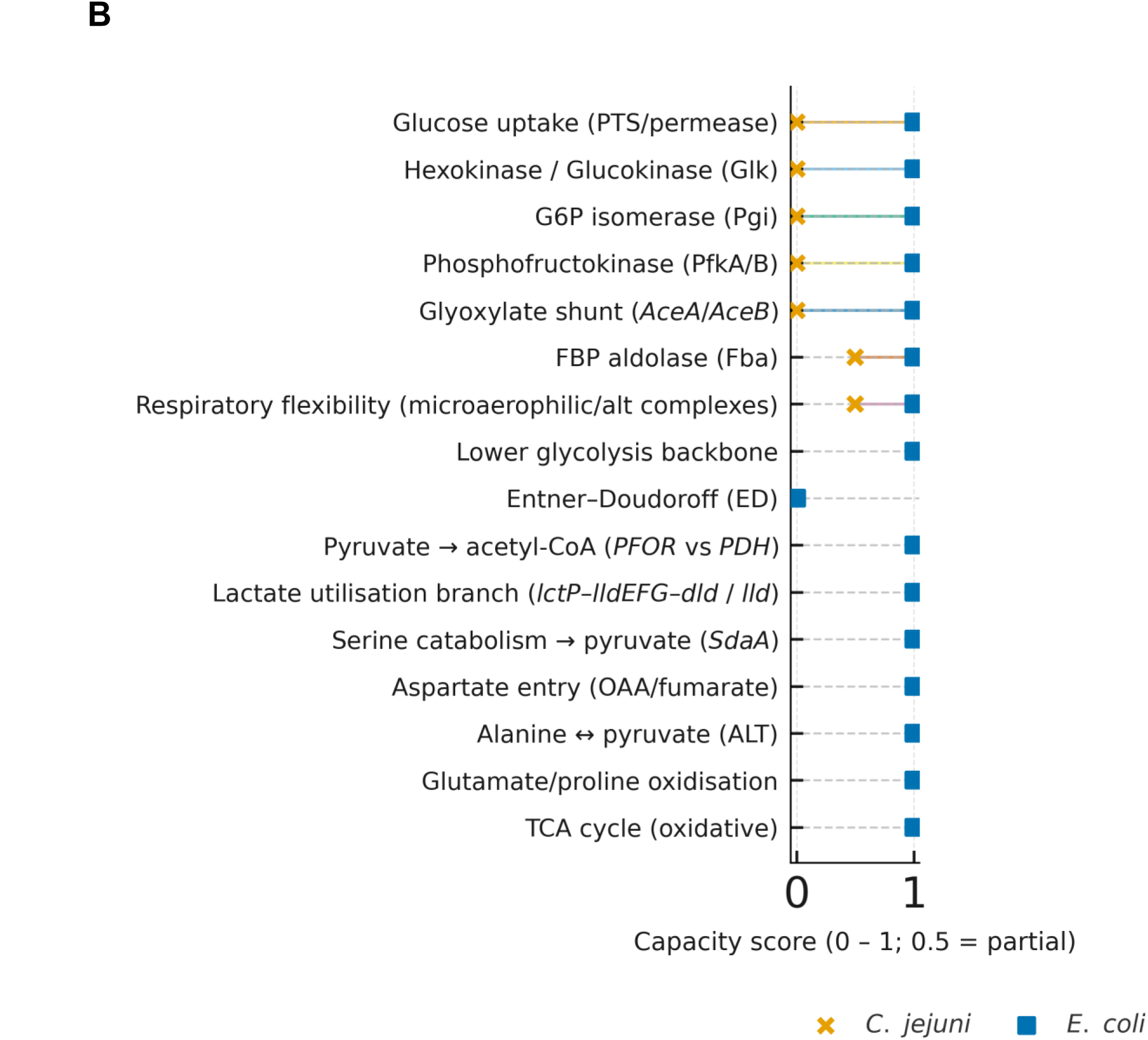
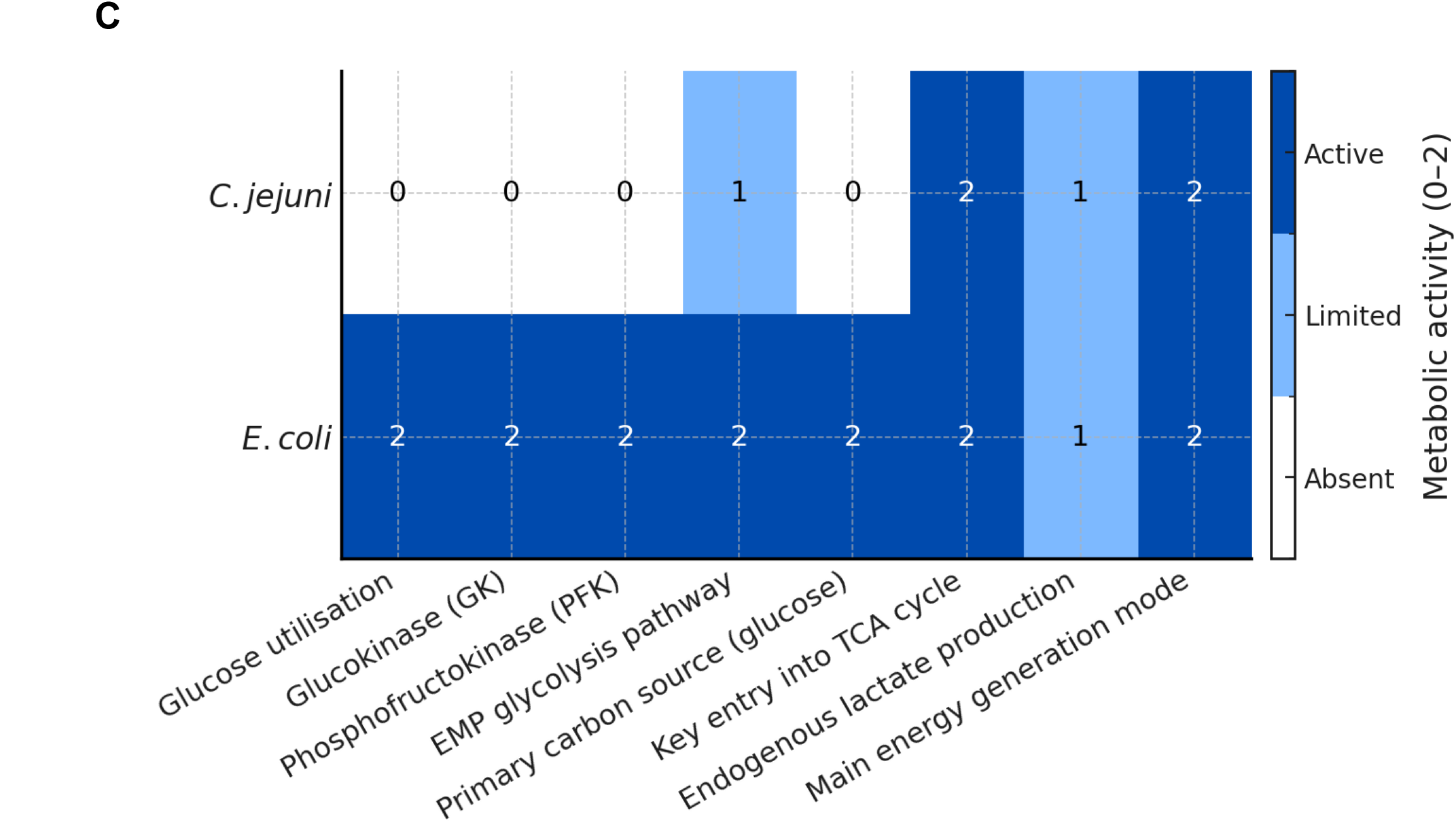
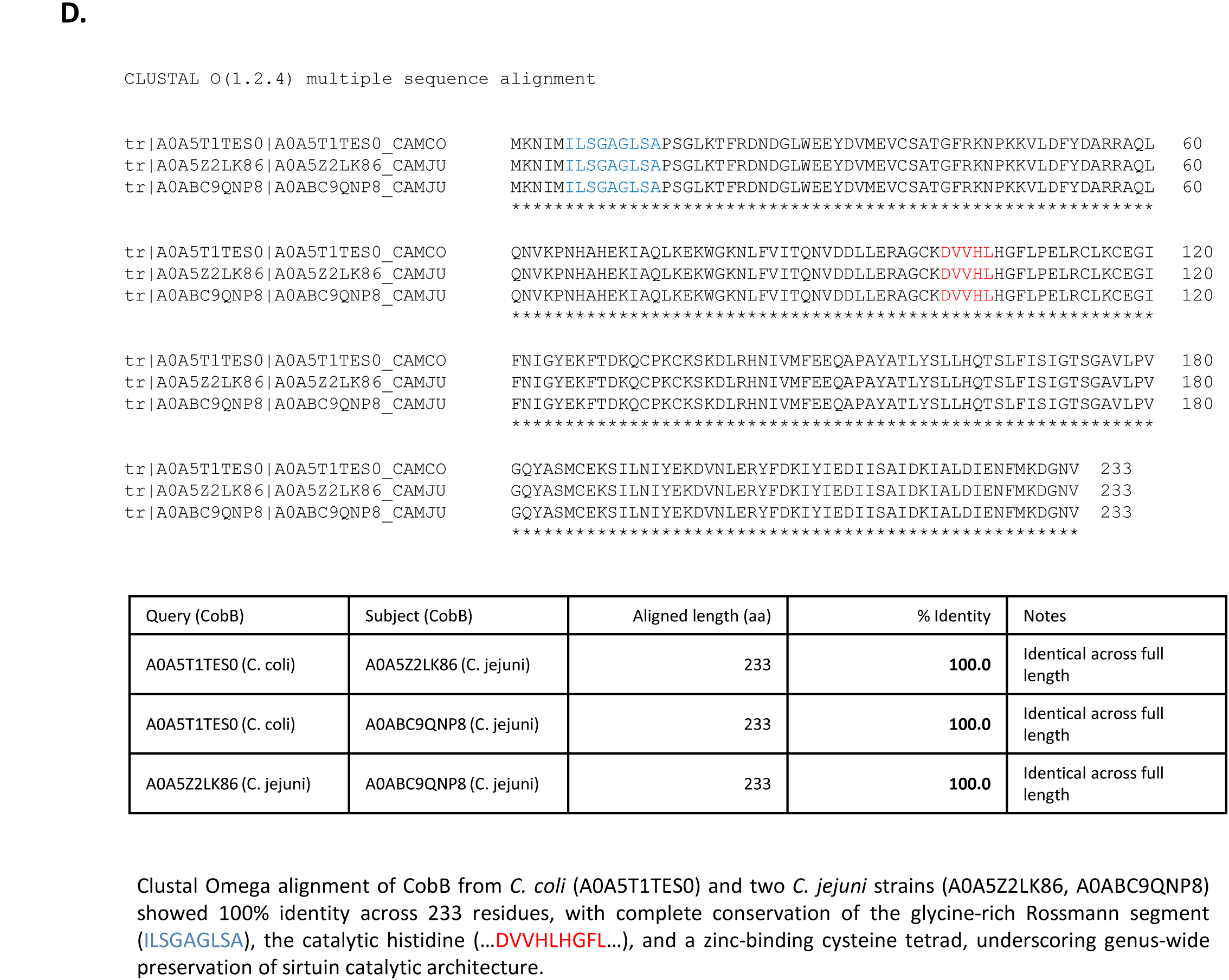
Amino acid–driven metabolism facilitates endogenous lactate generation and lysine lactylation in *Campylobacter jejuni*. **(A)** Core metabolic architecture under microaerophilic conditions. *C. jejuni* relies on amino-acid oxidation; glucose glycolysis via EMP is incomplete. Serine (SdaA) and alanine (AlaAT) feed the pyruvate node; aspartate enters the TCA via AspA (to fumarate) or AspAT (to oxaloacetate; OAA). Lactate is imported by LctP and oxidised in the periplasm by NAD-independent LDHs (LldEFG for L-lactate; Dld for D-lactate), with electrons transferred to the quinone pool and the cbb₃-type oxidase; the reverse pyruvate to lactate reaction is likely disfavoured *in vivo*. PFOR converts pyruvate to acetyl-CoA, which partitions to acetate (ATP-yielding), formate (PFL-like), or the partial TCA for biosynthesis and respiration. Proposed pools of lactyl-CoA and lactoyl-glutathione provide acyl donors for protein Kla, enabling a rapid, biomass-independent regulatory layer that links host-derived lactate to metabolic control. Strain-variable sugar catabolism is shown for context and is not a dominant carbon source. **(B)** Semi-quantitative capacity scores 0 = absent, 0.5 = partial/conditional, 1 = present/robust for central carbon and redox pathways in *C. jejuni* (×) and *E. coli* (▪). **(C)** Comparative metabolic heat map summarising pathway features that distinguish amino-acid– driven respiration in *C. jejuni* from glycolysis/fermentation in *E. coli*. **(D)** Table 2 showing metabolic contrasts between *C. jejuni* and *E. coli* (glycolytic model) under microaerophilic conditions. Amino-acid catabolism dominates carbon entry in *C. jejuni*, whereas *E. coli* primarily uses carbohydrate glycolysis. Respiratory, periplasmic lactate oxidation in *C. jejuni* links host-derived lactate to central metabolism and Kla.

**Table 1.** Key resources table Strains, Media, Antibodies Used in This Study.

| Reagent / Material | Source | Reference | Notes |
| --- | --- | --- | --- |
| *Campylobacter jejuni* NCTC 11168H | LSHTM *Campylobacter* collection | (Karlyshev *et al.*, 2002) | Reference wild-type strain |
| *Campylobacter jejuni* 81-176 | LSHTM *Campylobacter* collection | (Korlath *et al.*, 1985) | Reference wild-type strain |
| *Campylobacter coli* M8 (Chicken isolate) | This study | (Ghosh *et al.*, 2025) | Newly isolated wild-type strain |
| Columbia Blood Agar (CBA) | Oxoid | CM0331 | Routine culture medium |
| Brucella Broth (BB) | Oxoid | CM0169 | Liquid culture medium |
| Sodium L-lactate (≥98%) | Sigma-Aldrich | L7022 | Lactate pulsing experiments |
| Tris–RIPA buffer components | Thermo Fisher Scientific | 89900 | Protein extraction |
| Protease inhibitor cocktail | Roche | 11836153001 | Added during lysis |
| Qubit™ Protein Assay Kit (Broad Range) | Thermo Fisher Scientific | Q33211 | Protein quantification |
| 4-12% SDS–PAGE reagents and gels | Invitrogen | NP0315BOX | Protein separation |
| iBlot 3 Dry Blotting System | Invitrogen | IB33001 | Membrane transfer |
| Rabbit polyclonal anti–lactyl-lysine | Fisher | 17219403 | Primary antibody |
| IRDye 800CW anti-rabbit IgG | LI-COR Biosciences | 926-32211 | Secondary antibody |
| Odyssey M Imaging System | LI-COR Biosciences | Model M | Near-infrared detection |
| Image Studio Lite (v5.2) | LI-COR Biosciences | Free software | Band quantification |
| FIJI (ImageJ v1.54) | NIH | Open source | Image processing |
| BioRender | BioRender<br>(https://BioRender.com/0r3gzup) |  | Fig.1A Schematic generation |

**Table 2.** Comparative pathway matrix—amino acid–driven respiration in *C. jejuni* vs glycolysis in *E. coli*.

| Pathway / Feature | *C. jejuni* | *E. coli* (glycolytic model) | Functional implication |
| --- | --- | --- | --- |
| **Primary carbon source** | Amino acids (serine, alanine, aspartate; also glutamate/proline) | Glucose and other carbohydrates | *C. jejuni* derives carbon/energy mainly from amino-acid oxidation. |
| **Glycolytic capacity (EMP)** | **Incomplete**; lacks **pfkA** (PFK) and **pyk** (pyruvate kinase); sugar use is strain-variable | **Complete** EMP glycolysis | Glycolysis is largely inactive in *C. jejuni*; pyruvate comes predominantly from amino acids. |
| **Serine catabolism** | L-Ser → pyruvate (**SdaA**; high-flux) | Serine → pyruvate (secondary route) | Major pyruvate source in *C. jejuni*; provides carbon intersecting lactate pathways. |
| **Aspartate catabolism** | Asp → fumarate (**AspA**) and/or via **AspAT** to OAA → TCA | Asp feeds TCA | Supports oxidative metabolism and fumarate respiration at low O<sub>2</sub>. |
| **Pyruvate conversion** | Pyruvate → **acetyl-CoA** via **PFOR**; **no canonical NAD-LDH**; reduction to lactate is **not favored in vivo** | Pyruvate → acetyl-CoA (PDH) or → lactate (LDH) | *C. jejuni* channels pyruvate to acetyl-CoA; lactate production is minimal. |
| **TCA cycle** | Operational but **branched** under microaerobiosis | Complete and flexible (aerobic/anaerobic) | Coupled to amino-acid oxidation rather than sugar catabolism in *C. jejuni*. |
| **Energy generation (ATP)** | **Oxidative phosphorylation** linked to amino-acid and organic-acid oxidation; limited SLP (acetate kinase) | SLP and oxidative phosphorylation | *C. jejuni* relies mainly on respiration-linked ATP yield. |
| **Redox homeostasis** | Quinone-linked periplasmic dehydrogenases; **ferredoxin/ETC** cycling; **cbb<sub>3</sub>** oxidase | Balanced by glycolysis, fermentation, and ETC | Microaerophilic redox wiring sensitizes *C. jejuni* to lactate-linked electron flow. |
| **Lactate production** | Endogenous production **minimal**; reverse LDH activity disfavored; responses induced by exogenous lactate | Major fermentation product from carbohydrate metabolism | In *C. jejuni*, lactate mainly acts as an external substrate/signal. |
| **Lactate utilization** | **LctP** uptake; **NAD-independent LDHs** (**LldEFG/Dld**) oxidize lactate → pyruvate + e<sup>-</sup> → **Q** → **cbb<sub>3</sub>** | Consumed or produced depending on conditions | Couples host-derived lactate directly to energy metabolism. |
| **Key redox enzymes** | **PFOR**, **LldEFG/Dld**, **fumarate reductase**, **hydrogenase**, **cbb<sub>3</sub>** oxidase, **NDH-2/other dehydrogenases\*** | **PDH**, **LDH**, **NDH-1/2**, cytochrome **bo/bd** oxidases | Distinct respiratory architecture enables lactate sensing and supports Kla. |

1. Sugar uptake and upper EMP activity in *C. jejuni* are strain-dependent; table reflects canonical laboratory strains. 2. *C. jejuni* lacks a canonical NAD-dependent LDH; reverse pyruvate→lactate flux is **not favored** under tested conditions. 3. The set of respiratory dehydrogenases varies by strain; NDH-2/alternative dehydrogenases are listed as representative. 4. Proposed linkage to lysine lactylation (Kla) arises from lactyl-donor pools (lactyl-CoA and/or lactoyl-glutathione) generated during lactate metabolism; modification can increase without changes in biomass.

### Amino acid–driven metabolism facilitates *Campylobacter jejuni* for endogenous lactate generation and lysine lactylation

Under microaerophilic conditions, amino-acid oxidation sustains TCA cycle flux. L-serine is a dominant entry point: serine dehydratase (SdaA; *sdaA*) irreversibly converts serine to pyruvate, which is oxidised to acetyl-CoA by pyruvate:ferredoxin oxidoreductase (PFOR) or reduced to lactate by putative NADH-dependent oxidoreductases, whose precise identities remain to be defined in *Campylobacter*. This reduction regenerates NAD⁺ and supports measurable endogenous lactate formation in the absence of classical glycolysis.

A conserved *lctP–lldEFG–dld* locus (Fig. 3A) encodes an incomplete lactate uptake/oxidation system that couples host-derived lactate to intracellular redox balance. Within this architecture, lactate acts less as a terminal product and more as a redox-linked intermediate and potential signal, consistent with *C. jejuni*’s strictly respiratory lifestyle. Comparative network scoring (Fig. 3B) supports this specialisation: *E. coli* channels glucose through a complete EMP and fermentative apparatus and can efficiently re-assimilate lactate, whereas *C. jejuni* operates an amino acid–driven, respiration-linked programme in which lactate principally supports redox cycling rather than serving as a major carbon source. A pathway-feature Table 2 matrix and heat map (Fig. 3C) consolidate these differences and highlights *C. jejuni*’s commitment to amino acid catabolism in microaerophilic environments.

Integrating these observations, we propose that serine-derived pyruvate and the resulting lactate form flexible central nodes that couple carbon flux to post-translational control. Elevated lactate generated endogenously in amino acid–rich conditions or supplied exogenously in host niches such as the avian caecum and human colon (millimolar range) would favour formation of candidate acyl-donor intermediates, including lactyl-CoA or lactoyl-glutathione (proposed), enabling site-selective lysine lactylation on proteins within the ∼15–115 kDa range observed by immunoblotting (see Fig. 1–2). A bioinformatically identified CobB-like sirtuin (Fig. 3D) suggests potential reversibility; functional validation will require genetic and biochemical testing. Together, this framework suggests *C. jejuni* as an amino acid– respiring, redox-tuned pathogen that integrates central metabolism with post-translational regulation via lysine lactylation. In this context, lactate is not an endpoint but a signalling metabolite that coordinates adaptation and persistence in host-associated, microaerophilic niches (Fig. 3A–D).

### Proposed functional framework for lysine lactylation in *Campylobacter*

Unlike fermentative bacteria, *C. jejuni* relies on amino-acid oxidation to power respiration under low O₂. (Guccione *et al*., 2008, Hofreuter *et al*., 2008). Oxidation of serine, aspartate, and glutamate yields pyruvate, which either enters the TCA cycle or is reduced to L-lactate (Stahl *et al*., 2012, Luethy *et al*., 2017). Transient lactate pools arise endogenously (pyruvate reduction) and exogenously (host-derived uptake) (Mendz *et al*., 1997, Thomas *et al*., 2011). We propose that these pools seed formation of lactyl-CoA or lactoyl-glutathione, which act as activated donors for Kla on a defined set of cytoplasmic and periplasmic proteins (Supplementary Table 3).

In this model, periplasmic oxidation of L- and D-lactate by LldEFG/Dld transfers electrons to the quinone pool, sustaining the proton-motive force (Δp) (Fig. 4A). In the cytoplasm, lactyl-CoA is proposed to donate lactyl groups to lysine residues either non-enzymatically or via an acyltransferase, with CobB-mediated delactylation favored under oxidised conditions (high NAD⁺) (Fig. 4B). We propose Kla on periplasmic and membrane-associated proteins, including PglB, SodB, and KatA, suggesting impacts on glycan-precursor flux, secretion, and host interactions (Fig. 4C). These would be consistent with a redox-coupled mechanism in which reduced states (QH₂↑, NADH↑, NAD⁺↓) diminish CobB activity and elevate Kla, whereas oxidation reverses this (Fig. 4D). Accordingly, Kla may function as a readout of intracellular redox and a regulator of envelope remodeling in *Campylobacter*, with implications for virulence and fitness. The reproducible ∼42–45 kDa Kla-reactive band observed across *C. jejuni* 11168H, 81-176, and *C. coli* (Figs. 2–3) is consistent with a conserved lactylated enzyme operating at the pyruvate–lactate node. Proposed candidate targets (Table. 3) include enzymes linked to serine/pyruvate catabolism (e.g., SdaA, PorA, LldD) and periplasmic or surface-exposed proteins that encounter redox stress or host-derived lactate. Many of these proteins harbour solvent-exposed lysines amenable to enzymatic acylation or to non-enzymatic transfer under elevated intracellular lactate/acyl-CoA. The modification appears sub-stoichiometric yet reproducible features characteristic of regulatory PTMs rather than bulk acylation. Reversibility is plausibly mediated by CobB-like sirtuin delactylases, providing a mechanism to reset modification states under energy-limiting or oxidative stress conditions (Fig. 3D). Amino acid motif-guided, *in silico* site prediction flagged SodB (superoxide dismutase [Fe]) as the highest-scoring Kla candidate in this dataset, with a predicted lysine at K29 within the peptide TFSYHHGKHHNTYVT (motif position K29) returning top scores (motif score 15.9; conservation 7.22; classifier call: Very strong). Given SodB’s central role in detoxifying superoxide and maintaining redox poise, lactylation at K29 would place Kla directly at the redox–metabolism interface.

**Figure 4.**
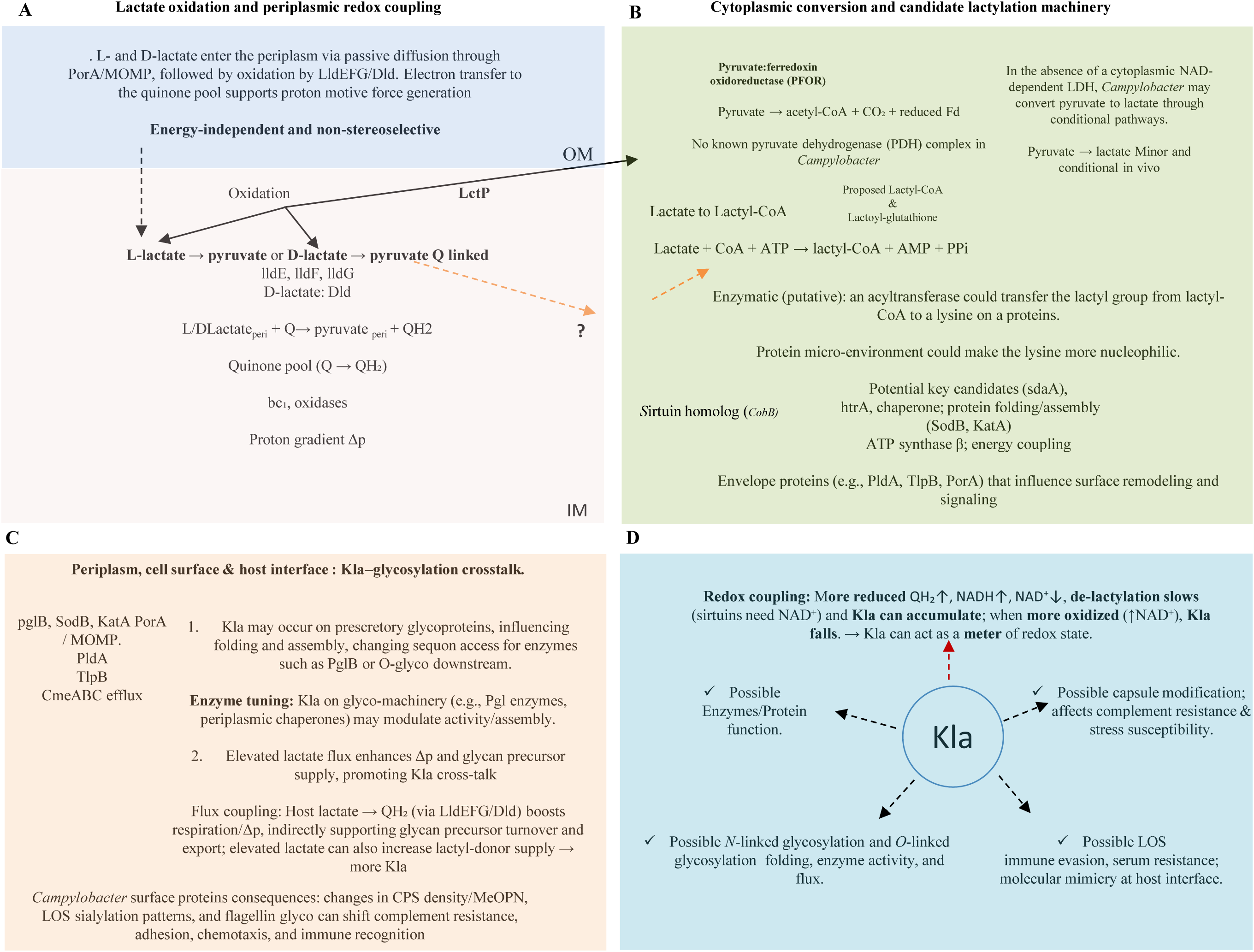
Conceptual framework for Kla as a metabolic signal in *Campylobacter*. (A & B) Host-derived lactate and respiratory coupling. Exogenous L- and D-lactate from the host–microbiota niche (chicken caecum ≈ 20–30 mM; human gut lumen up to ∼10 mM) are imported via *LctP*. In the periplasm, *LldEFG* (L-lactate) and *Dld* (D-lactate) oxidise lactate to pyruvate in an NAD-independent, membrane-linked reaction that reduces the quinone pool (Q → QH₂). QH₂ feeds the respiratory chain (cytochrome *bc₁*/alternative oxidases) to generate a proton motive force (Δp) under microaerophilic conditions. Pyruvate serves as a central node, supplying PorAB (PFOR) to acetyl-CoA and anaplerotic/biosynthetic pathways. A classical NAD-dependent LDH is absent; pyruvate to lactate reduction is thought to be minimal *in vivo* (dashed). We propose formation of lactyl-CoA via an acyl-CoA synthetase (lactate + CoA + ATP → lactyl-CoA + AMP + PPi), with possible non-enzymatic routes (grey, dashed). Lactyl-CoA (or related donors, e.g., lactoyl-glutathione) may drive protein lysine lactylation. Cytoplasmic module: redox coupling and reversibility. Site-selective Kla is proposed on enzymes such as SdaA, DnaK, and HtrA, tuning activity, complex assembly, or substrate affinity to couple NADH/NAD⁺ state to metabolism. Reversibility via an NAD⁺-dependent CobB-type sirtuin would establish a write–erase cycle for rapid adjustment to redox and flux changes. **(C)** Envelope module: host interface and Kla–glyco crosstalk. Periplasmic and membrane proteins, including PldA, TlpB, and PorA may acquire lactyl marks in response to host-derived lactate, linking metabolic state to envelope remodelling and signalling. Potentially glycosylated surface proteins (FlgE, FlgL, CmeC) represent potential nodes of Kla– glycosylation crosstalk with effects on stability, secretion, and immune recognition. **(D)** Integrated proposed model. Across compartments, Kla acts as a metabolic translator that integrates amino-acid catabolism, redox homeostasis, modification of the LOS and capsule, and virulence adaptation, positioning lactate as a signalling molecule rather than a terminal product in microaerophilic *Campylobacter*.

This framework yields a dual-compartment view in which Kla coordinates metabolic flux with host interaction: periplasmic oxidation of L- and D-lactate by LldEFG/Dld feeds the quinone pool, while cytoplasm formation of lactyl donors (putatively lactyl-CoA/lactoyl-glutathione, via an acyl-CoA synthetase–type route) supports site-selective lactylation of redox/energy proteins (e.g., PFOR/PorAB, SdaA, AtpD, SodB, KatA). In parallel, presecretory forms of envelope proteins (e.g., CmeA/CmeC, PorA/MOMP, FlgE/FlgL) can be lactylated in the cytoplasm and subsequently displayed in the periplasm/outer membrane, where Kla may modulate folding, export efficiency, or glycan display, establishing Kla–glycosylation crosstalk that links metabolism to envelope physiology. Reversible delactylation by CobB-type sirtuins would confer rapid responsiveness to oxygen tension and host-derived metabolites, forming a redox-linked post-translational control circuit.

## Discussion

Since the pioneering work of Skirrow and colleagues in the late 1970s, understanding of the molecular basis of *Campylobacter jejuni* virulence and persistence has advanced significantly, yet many key features remain unresolved (Skirrow, 1977, Blaser *et al*., 1983). Despite a compact genome of ∼1.6 Mb, with notable plasticity, a paucity of canonical two-component regulators, and heavy reliance on phase variation, *C. jejuni* displays striking physiological plasticity and efficient colonisation reaching ∼10⁹ CFU g⁻¹ in avian caeca and establishing human infection with an inoculum as low as ∼500 cells. (Robinson, 1981; Beery *et al*., 1988; Black *et al*., 1988). Unusually for an enteric pathogen, *C. jejuni* encodes both *N-*linked and *O-*linked post-translational protein glycosylation systems that contribute to multiple phenotypes (Goon *et al*., 2003, Szymanski *et al*., 2003, Linton *et al*., 2005, Ewing *et al*., 2009). The co-existence of these glycosylation pathways with Kla reported here points to a richer post-translational landscape than previously appreciated one capable of integrating environmental cues with adaptive physiology.

Our data support a model in which lactate abundant in the avian caecum and present in the human intestine act as a redox-sensitive signalling metabolite. In *C. jejuni*, the EMP pathway is incomplete; central carbon flux is supplied by amino-acid oxidation, creating a metabolic architecture in which transient pools of L-lactate arise both endogenously and via uptake (Parkhill *et al*., 2000, Hofreuter, 2014). The partially characterised lactate locus *cj0076c– cj0075c–cj0074c–cj0073c* comprises a lactate transporter (*lctP*, *cj0076c*) and a non-flavin, iron–sulphur, three-subunit membrane oxidoreductase (*cj0075c–cj0074c–cj0073c*) that oxidises L-lactate to pyruvate (Thomas *et al*., 2011, Sinha *et al*., 2024). We propose that these lactate pools seed formation of intracellular lactyl-CoA or lactoyl-glutathione, which then act as acyl donors for Kla on a defined set of cytoplasmic, membrane, and periplasmic proteins. The robust, reproducible ∼42–45 kDa band observed across *C. jejuni* 11168H, 81-176, and *C. coli* M8 including in the absence of exogenous lactate argues for a conserved, constitutively lactylated target at the pyruvate–lactate node. Responsiveness to L-lactate pulses indicates stereoselective control, consistent with an enzymatic step that recognises the L-enantiomer rather than indiscriminate non-enzymatic acylation.

Mechanistically, we hypothesise that in *Campylobacter*, a streamlined transcriptional programme is complemented by Kla as a compensatory post-translational layer. By modulating catalytic efficiency, substrate affinity, or protein–protein interactions, Kla could couple intracellular redox state (e.g., NADH/NAD⁺ and quinone-pool status from periplasmic, NAD-independent lactate oxidation) to enzyme activity and pathway routing around the PFOR-centered pyruvate node. The observed selectivity and sub-stoichiometry, Kla bands in Figures 2 and 3 that appear only at higher protein loads, are hallmarks of a regulated PTM rather than bulk, non-specific acylation, and are consistent with a model in which lactyl-donor availability tunes flux through key *Campylobacter* pathways. Candidate targets (Supplementary Fig. 5) include enzymes integral to serine/pyruvate catabolism and redox control (SdaA, PorA, LldD, AspA, Pgk, PglK, SodB, and components of the PstSCAB system) as well as surface-exposed or envelope-associated proteins (CmeC, MOMP, FlgE, FlaA, FlpA, Peb1A, Peb4, JlpA) that encounter host-derived metabolites and redox stress. Reversibility is possibly provided by CobB-like sirtuin delactylases, establishing a write–erase cycle that aligns metabolic state with proteome function. Definitive evidence for enzyme-mediated lactyl transfer and delactylation will require genetic and biochemical dissection of candidate acyltransferases and sirtuins, together with stereochemical controls using D-lactate, although *C. jejuni can also use D-lactate*.

*In silico* mapping of putative lactylation sites in *Campylobacter* (Table 3) identified conserved sequence contexts and solvent-exposed lysines across metabolic and surface-associated proteins. These predictions were guided by acyl-lysine amino acid motifs reported for eukaryotic acyltransferases (e.g., EP300/CREBBP, KAT families) and are presented as hypothetical bacterial acyltransferases that remain to be determined experimentally (Deng *et al*., 2016). A good candidate emerging from this analysis is SodB (*sodB*; Cj0169c), the Fe-dependent superoxide dismutase that anchors cytoplasmic oxidative-stress control (Purdy *et al*., 1999). We identified a very strong Kla prediction at K29 (peptide TFSYHHGKHHNTYVT; motif score 15.9; conservation 7.22), the highest-scoring site in our dataset. A lactyl mark at K29 is well positioned to influence metal coordination, oligomer stability, or partner interactions, thereby coupling NADH/NAD⁺ balance and lactate availability to antioxidant capacity. Within our framework, SodB provides a direct mechanistic bridge between amino-acid-driven redox metabolism and proteome regulation: when lactate pools rise, Kla-dependent modulation of SodB could recalibrate reactive-oxygen-species buffering without wholesale transcriptional responses.

To pre-empt over-interpretation, SodB K29 remains a prediction pending direct mass spectrometry-based validation. A concrete path forward includes: (i) Kla immuno-enrichment followed by PRM/SRM targeting the TFSYHHGKHHNTYVT peptide; (ii) site-directed mutagenesis (K29R non-acylatable; K29Q/E lactyl-mimetic) and (iii) CobB-dependent delactylation tests to assess reversibility. Positive results would establish SodB as a nodal Kla substrate linking carbon flux, redox homeostasis, and stress fitness.

Ecologically, positioning lactate as both cue and co-substrate reconciles *C. jejuni*’s strict respiratory lifestyle with success in oxygen-limited niches. Rather than fully reducing pyruvate to lactate for fermentation, *C. jejuni* appears to maintain lactate as a transient redox intermediate that conserves reducing power, sustains respiratory electron flow most plausibly via quinone-linked entry into the electron-transport chain and furnishes acyl donors for post-translational control. This interface between host/microbiota metabolism and bacterial regulation suggests translational strategies: perturbing lactate flux or stereochemistry, biasing microbiome-derived metabolite availability, or targeting Kla cycling to attenuate colonisation and persistence. Such approaches could complement antimicrobial stewardship in settings where *Campylobacter* is prevalent and may, pending validation, extend to other microaerophilic, phylogenetically related epsilon-proteobacterial pathogens, such as *Helicobacter pylori* and *Arcobacter* species where the functional relevance of lysine lactylation remain to be established. Additionally, the ε-proteobacteria subdivision of bacteria often inhabit harsh environments such as deep-sea vents (Huber *et al*., 2007) and frequently have the *N-*linked post-translational glycosylation systems (Mills *et al*., 2016) suggesting post-translational modification is a common feature of this sub-division of bacteria.

In conclusion, our data support a model in which *C. jejuni* uses lactate as a signal that modulates its proteome via lysine lactylation. This finding reframes lactate metabolism as a regulatory bridge linking carbon flux, redox homeostasis, and the glycan-rich envelope. Further research could involve (i) delineating the enzymatic machinery and stereochemistry of lactyl-donor formation; (ii) mapping Kla sites and stoichiometry across growth states, gut-mimetic media (A.E manuscript in preparation), and oxygen tensions, and examining links to kin recognition and non-kin discrimination in *Campylobacter* (A.E manuscript in preparation); (iii) defining functional consequences for specific targets, including the conserved ∼42–45 kDa protein; and (iv) testing whether CobB-type delactylases reset signalling during oxidative or nutrient stress. More broadly, Kla may constitute a general mechanism by which environmental metabolites shape bacterial behaviour and possibly pathogenesis.

## Author Contributions

**A.E. (Abdi Elmi):** Conceptualisation; Study design; Methodology; Investigation; Validation; Formal analysis; Visualisation; Writing – original draft; Writing – review & editing; Project administration. **B.W. (Brendan Wren):** review & editing.

## Author statement

A.E. conceived the project independently, managed, performed all experiments, analysed the data, prepared figures and legends, and wrote and revised the manuscript.

## Acknowledgments

We thank **Joseph Moulton** for technical support and data collection, and **Dr Richard Stabler** for providing *Campylobacter coli* strain M8.

## Inclusion & Diversity Statement

This project was conceived, executed, and written independently by the lead author (A.E.). We are committed to equitable credit and transparent authorship practices that recognise leadership by researchers from racially minoritised communities and other under-represented groups.

