## Supplemental Table 3 for "Lactate-responsive lysine lactylation in *Campylobacter* revealed by antibody-based detection of a conserved post-translational modification"

Supplementary Table S3. In silico mapping of predicted lysine lactylation sites across representative *Campylobacter jejuni* proteins

| **Protein (Gene ID)** | **Function / Localization** | **Representative Peptides (Position)** | **Predicted HAT** | **Highest Score** | **Cutoff** | **Predicted Status** | **Remarks / Functional Note** |
|---|---|---|---|---|---|---|---|
| **SdaA (Cj0828c)** | Serine dehydratase, cytoplasmic (PLP-dependent) | AELEKKGKNSNQNKK (159), SNQNKKKKLDIELNN (168), GSLSLTGKGHLSDKA (55) | CREBBP, KAT2B, KAT8 | **7.7** | 7.22 | ✅ Strong | Clustered lysines near catalytic pocket; high surface exposure and metabolic linkage |
| **PorA (Cj0115)** | Major outer membrane porin | ANHLGGGKKLEAVAR (374), NNSKQDHKYRAQVNF (65) | KAT8 | **10.2** | 7.22 | ✅ Strong | Surface-exposed β-barrel loops; potential role in host interaction |
| **CmeC (Cj0369)** | Outer membrane efflux channel | EDTSKNIKEESKNLD (480), YENENALKEAYKSAK (196) | CREBBP, KAT2A | **3.86** | 1.35 | ✅ Moderate–Strong | Extracellular loop lactylation; possible function in envelope stress adaptation |
| **AspB (Cj0525)** | Aspartate aminotransferase, cytoplasmic | ANFIKNYK (389), KIEKDSMKFCQKLLE (338) | CREBBP, KAT2A | **2.65** | 1.38 | ✅ Strong | Catalytic-domain lysines; potential redox-regulated modification |
| **TrxB (Cj0205)** | Thioredoxin reductase, cytoplasmic | GGKTELAKAVIVCTG (105) | KAT2B | **2.34** | 1.34 | ✅ Strong | Redox enzyme; consistent with lactylation–redox coupling model |
| **LysS (Cj0339)** | Lysine–tRNA ligase, cytoplasmic | RPLKSELKEKE (498), LKSELKEKE (500) | CREBBP | **4.25** | 1.35 | ✅ Strong | Lysine-rich catalytic domain; potential translational |

| | | | | | | | |
|---|---|---|---|---|---|---|---|
| | | | | | | | regulation node |
| **CjaA (Cj0982c)** | Amino acid transporter (periplasmic binding) | MKKILLSVL (2–3), KKGDKELKEFIDNLI (235) | KAT2B | **2.09** | 1.34 | ✅ Moderate | Signal-peptide and binding region; may alter transport or affinity |
| **FlgE (Cj1729)** | Flagellar hook protein, periplasmic | AQIITAHKYIYSSNP (444) | KAT8 | **7.8** | 7.22 | ✅ Strong | Periplasmic hinge region; possible modulation of flagellar motility |
| **Cbf2 (Cj0143)** | Peptidyl-prolyl cis-trans isomerase, periplasmic | NSAKVEYK (273), MKKFSLVAA (2–3) | CREBBP, KAT2B | **3.01** | 1.34 | ✅ Strong | Protein-folding chaperone; modification may influence OMP assembly |
| **FrdC (Cj0491)** | Fumarate reductase subunit, membrane | LGKSIEGKKSKMPAK (16) | KAT2A | **1.78** | 1.38 | ✅ Moderate | Electron transport component; candidate for redox-regulated lactylation |
| **GmhD (Cj1427c)** | GDP-D-heptose dehydrogenase, cytoplasmic | MSKKVLITGG (3–4) | KAT2B, KAT8 | **7.6** | 7.22 | ✅ Strong | Links LPS biosynthesis and metabolism; potential glycan–redox interface |
| **FlaA (Cj1339c)** | Flagellin A, surface | NVAALNAKANADLNS (15) | EP300 | **0.69** | 0.42 | ✅ Weak | Low-confidence site; potential exposure on filament surface |
| **CheA (Cj0019c)** | Chemotaxis histidine kinase, cytoplasmic | DQARRAQKKQTTNAA (195), QARRAQKKQTTNAAP (196) | KAT2A | **1.83** | 1.38 | ✅ Strong | Two-component regulator; may link |

| | | | | | | | |
|---|---|---|---|---|---|---|---|
| | | | | | | | redox and chemotaxis signaling |
| **GroEL (Cj1220)** | Chaperonin, cytoplasmic | AATETEMKEKKDRVD (389) | CREBBP | **1.88** | 1.35 | ✅ Strong | Folding chaperone; potential stress-linked lactylation |
| **AhpC (Cj0334)** | Alkyl hydroperoxide reductase, cytoplasmic | GVAEYLGKNEAKL (193) | CREBBP | **2.25** | 1.35 | ✅ Strong | Antioxidant enzyme; modification near catalytic site |
| **PglB (Cj1124c)** | Glycosyltransferase, inner membrane | DAKVFKLKI (712) | CREBBP | **2.13** | 1.35 | ✅ Strong | N-glycosylation enzyme; connects metabolism to protein glycosylation |
| **CadF (Cj1478c)** | Fibronectin-binding protein, outer membrane | GSRAYNQKLSERRAK (263) | KAT2A | **1.64** | 1.38 | ✅ Strong | Adhesion factor; integrates host interaction and metabolic adaptation |
| **CiaB (Cj0912)** | Invasion antigen B, secreted | SGEFERYKKK (608), EFERYKKK (610) | CREBBP | **3.07** | 1.35 | ✅ Strong | Secreted effector; lactylation may influence invasion signaling |
| **OorA (Cj0535)** | 2-Oxoglutarate:acceptor oxidoreductase A, cytoplasmic | TPSEIIAKVKENI (369) | CREBBP | **1.98** | 1.35 | ✅ Strong | Central metabolic enzyme; redox-modulated flux control candidate |
| **OorB (Cj0536)** | 2-Oxoglutarate:acceptor oxidoreductase B, cytoplasmic | EEVRRAAKEKRMVDL (270) | CREBBP | **2.33** | 1.35 | ✅ Strong | Electron-transfer enzyme; predicted metabolic redox target |
| **SodB** | Superoxide | TFSYHHGKHH | KAT8 | **15.9** | 7.22 | ✅ | Antioxidant |

| | | | | | | | |
|---|---|---|---|---|---|---|---|
| **(Cj0169c)** | dismutase [Fe], cytoplasmic | NTYVT (29) | | | | Very Strong | enzyme; highest-scoring lactylation site identified |
| **CheW (Cj0020)** | Chemotaxis coupling protein, cytoplasmic | ENILTILKVEALLKR (164) | CREBBP | **1.49** | 1.35 | ✅ Strong | Signal relay factor; potential chemotaxis regulation via lactylation |
| **Peb1A (Cj0921)** | Cell-binding factor, periplasmic | EIDALAKKWGL (256), EGKLESIKSKGQLIV (35) | CREBBP, KAT2B | **1.84** | 1.34 | ✅ Strong | Adhesion/transport factor; surface adaptation site |

*In silico mapping of lysine lactylation hotspots across representative cytoplasmic, membrane, and surface-associated proteins of* Campylobacter jejuni *NCTC 11168. Prediction scores were derived from GPS-Lactylation v1.0 and DeepKla (2024) algorithms using conserved EP300/CREBBP- and KAT-family acyltransferase recognition motifs from eukaryotic systems. Proteins were prioritized based on functional relevance, localization, and peptide context within catalytic or surface-exposed domains. Predicted lysine sites showing scores above tool-specific thresholds are classified as strong or moderate candidates for experimental validation.*
